# Gram-scale one-pot enzymatic synthesis of CDP-ribitol by a designed bifunctional fusion enzyme

**DOI:** 10.64898/2026.07.27.741123

**Authors:** Xuyang Zhang, Li Pan, Yumeng Wang, Xianwei Liu, Mingxia Bi, Peixue Ling, Congcong Chen, Shuaishuai Wang

## Abstract

D-Ribitol-5-phosphate (Rbo5P) plays vital roles in bacterial and mammalian development. In mammals, Rbo5P is an essential component of the O-mannosyl glycan on α-dystroglycan, and its defective biosynthesis causes a group of dystroglycanopathies. Supplementation with cytidine diphosphate ribitol (CDP-ribitol, CDP-Rbo) and its analogues has shown therapeutic potential for certain dystroglycanopathies. However, facile and scalable strategies to prepare CDP-Rbo remain unavailable. Here, we report a practical one-pot enzymatic strategy for gram-scale CDP-Rbo production. Starting from inexpensive ribitol, Rbo5P is first generated by L-ribulokinase (AraB) and then converted to CDP-Rbo by CDP-ribitol pyrophosphorylase (TarI) in 90% yield. To streamline the process, AraB and TarI were fused into a single bifunctional biocatalyst, AraB-TarI, which converts ribitol directly into CDP-Rbo on a gram-scale in a one-step reaction. Furthermore, a nucleotide recycling strategy was developed to lower the cost and increase the atom economy from 47% to 75%. This work provides a green and scalable route to CDP-Rbo and a reliable material supply for developing therapeutics against dystroglycanopathies.

## Introduction

α-Dystroglycan (α-DG) is a heavily glycosylated cell-surface protein that bridges the extracellular matrix and the intracellular cytoskeleton. This bridging role depends on the O-mannosyl glycosylation of α-DG. Mutations that disrupt this glycosylation cause a group of congenital muscular dystrophies (CMD), collectively termed dystroglycanopathies.^1^ The O-mannosyl glycan carries tandem D-ribitol-5-phosphate (Rbo5P) units that are required for its formation (Fig. 1a). Mutations in the isoprenoid synthase domain-containing protein (ISPD), which is involved in CDP-Rbo biosynthesis, cause a severe form of CMD.^2-4^

**Fig. 1.**
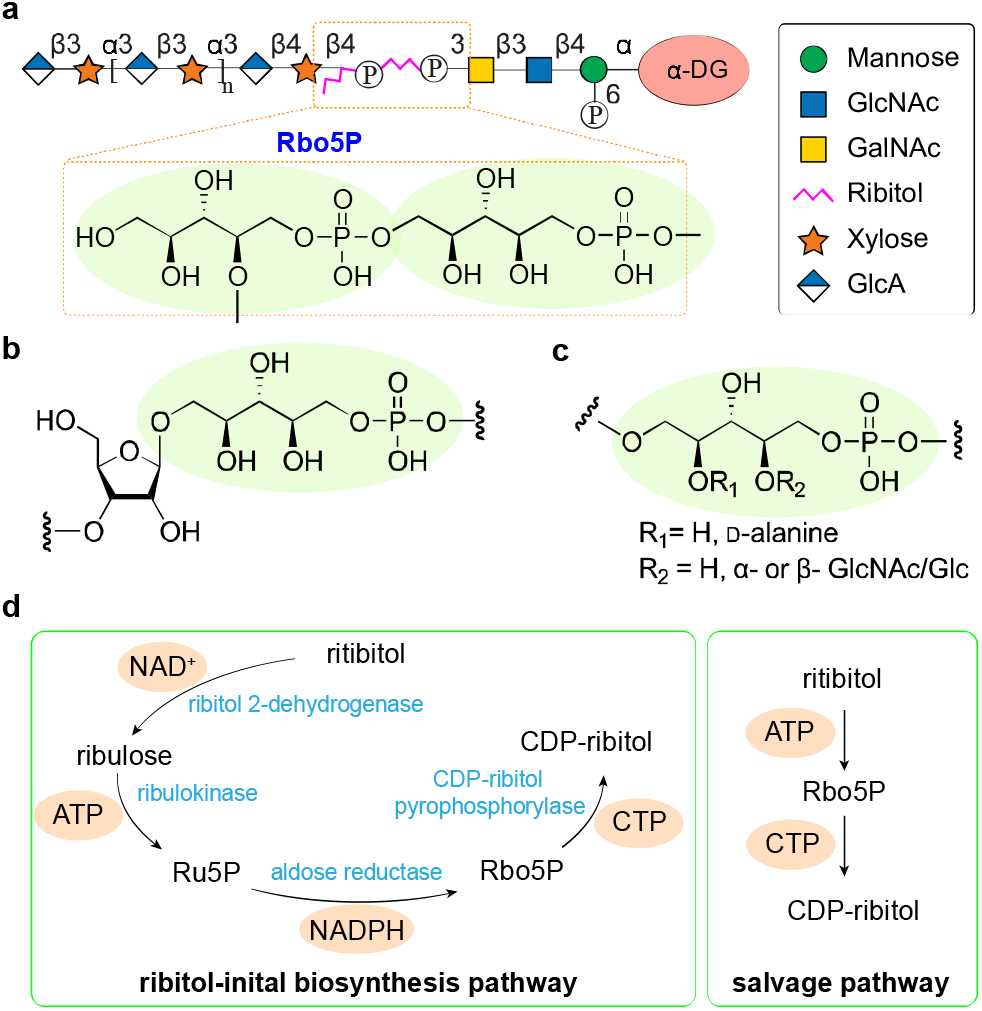
The Rbo5P structure in mammals and bacteria and the biosynthesis pathway of CDP-Rbo. (a) The tandem Rbo5P in O-mannosyl glycan. (b) Rbo5P in the capsular polysaccharide of Gram-negative pathogens Hib. (c) Rbo5P in the WTA of Gram-positive pathogens *Staphylococcus aureus*. (d) The ribitol-initiated de novo biosynthesis pathway and salvage pathway for the synthesis of CDP-Rbo.

Rbo5P is not restricted to mammals, it is also a key component of the bacterial cell wall. It occurs in the wall teichoic acid (WTA) of the Gram-positive pathogen *Staphylococcus aureus*^5-7^ and in the capsular polysaccharide of the Gram-negative pathogen *Haemophilus influenzae* type b (Hib) (Fig. 1b–c).^8^ These Rbo5P-containing polymers serve as virulence factors that help the pathogens to evade the host immune system.

Cytidine diphosphate D-ribitol (CDP-ribitol, CDP-Rbo) is the active form of Rbo5P, which is utilized by glycosyltransferases to generate Rbo5P-containing glycans. Recently, supplementation with CDP-Rbo or its analogues has been shown to be beneficial in ISPD-deficient muscular dystrophy, which provides a promising therapeutic approach as no treatment for ISPD-deficient muscular dystrophy is currently available.^9^

Due to the significant biological functions and great potential of CDP-Rbo for biomedical applications, several approaches have been reported for its preparation. CDP-Rbo was first isolated from *Lactobacillus arabinosus*.^10^ Then, a chemical approach was reported for the synthesis of CDP-Rbo.^11^ Later, different chemical strategies were developed to access a range of CDP-Rbo derivatives (Fig. 2a).^12, 13^ Enzymatically, CDP-Rbo has been prepared from D-ribulose 5-phosphate (Ru5P) using the bifunctional Bcs1 from *H. influenzae*,^14^ or Rbo5P using TarI (Fig. 2b).^15, 16^ However, a practical strategy for large-scale synthesis of CDP-Rbo is still lacking as current routes require costly phosphorylated substrates (Rbo5P, Ru5P, or ribose 5-phosphate) and cofactors (NAD^+^, NADPH).

**Fig. 2.**
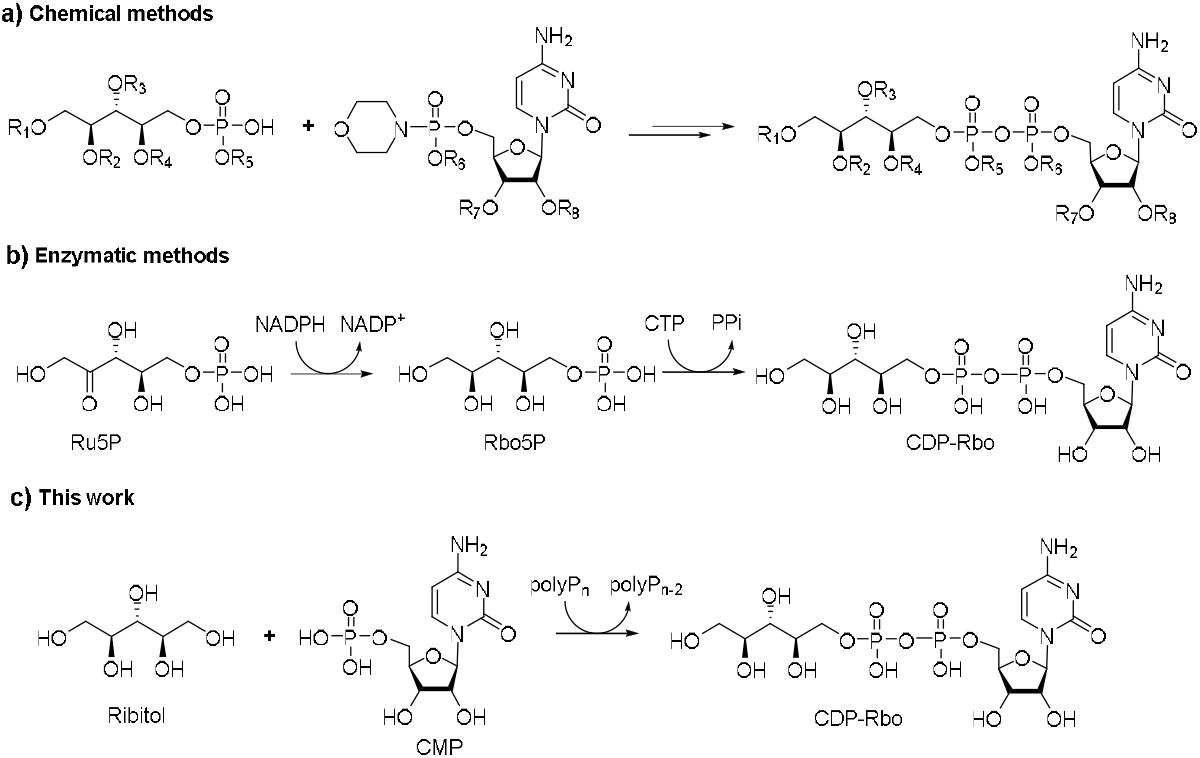
Strategies for the preparation of CDP-Rbo. a. Chemical methods for the synthesis of CDP-Rbo analogues; b. Enzymatic methods for the generation of CDP-Rbo by starting with D-ribulose 5-P (Ru5P). c. The enzymatic methods developed in this work starting with ribitol, CMP and poylPn.

In the biosynthesis of sugar nucleotides, two routes are generally available: a *de novo* pathway and a salvage pathway. For *de novo* synthesis of CDP-Rbo, D-ribitol is first oxidized to D-ribulose and then phosphorylated to Ru5P, an intermediate also shared with the pentose phosphate pathway (Fig. 1d)^17^. The reduction of Ru5P by NADPH yields Rbo5P, which is then cytidylylated to yield CDP-Rbo.^18-22^ A four-enzyme cascade with four cofactors is required for this pathway. By contrast, the salvage pathway requires only two enzymes and two cofactors without consuming NAD^+^ or NADPH. This lower cofactor cost makes it more efficient than the *de novo* pathway. However, the enzymes responsible for the salvage pathway have yet to be identified, although several candidates have been proposed.^23-25^

Here, we report a practical and sustainable enzymatic strategy to produce CDP-Rbo from low-cost and readily available ribitol (Fig. 2c). Two enzymes were employed for the generation of CDP-Rbo to mimic the salvage pathway. Ribitol was first phosphorylated to Rbo5P by L-ribulokinase (AraB) in an ATP-dependent reaction, and Rbo5P was then coupled with CTP by CDP-Rbo pyrophosphorylase (TarI) to give CDP-Rbo. To simplify the process, we fused the two enzymes into a bifunctional CDP-Rbo synthase (AraB-TarI) that converts ribitol directly into CDP-Rbo in a one-pot reaction on a gram scale. Furthermore, a nucleotide recycling strategy was employed to replace the stoichiometric consumption of ATP and CTP with the cheaper AMP and CMP. This approach provides efficient and economical access to CDP-Rbo and supplies a key material for developing therapeutics targeting ISPD-deficient dystroglycanopathies.

## Results and discussion

### Enzymatic phosphorylation of ribitol by AraB

The FGGY family comprises ATP-dependent kinases that phosphorylate a range of sugar alcohols and monosaccharides, such as erythritol, glycerol, ribulose and xylulose, and its members often display broad substrate specificity.^26^ As no native kinase has been reported for the phosphorylation of ribitol to Rbo5P, we searched this family for an enzyme with ribitol kinase activity. Reported FGGY kinases include L-ribulokinase from *E. coli*^27, 28^ and D-ribulokinases from *E. coli*,^29^ *Klebsiella aerogenes*,^30, 31^ *K. pneumoniae*,^17^ *Saccharomyces cerevisiae* and human.^23^ Because D-ribulokinase has been proposed to have evolved from a subgroup of L-ribulokinase with more restricted substrate specificity,^26^ we reasoned that a broad-specificity L-ribulokinase might retain promiscuous activity toward D-ribitol. We therefore selected L-ribulokinase (AraB) from *E. coli* K-12 (Fig. 3a).

**Fig. 3.**
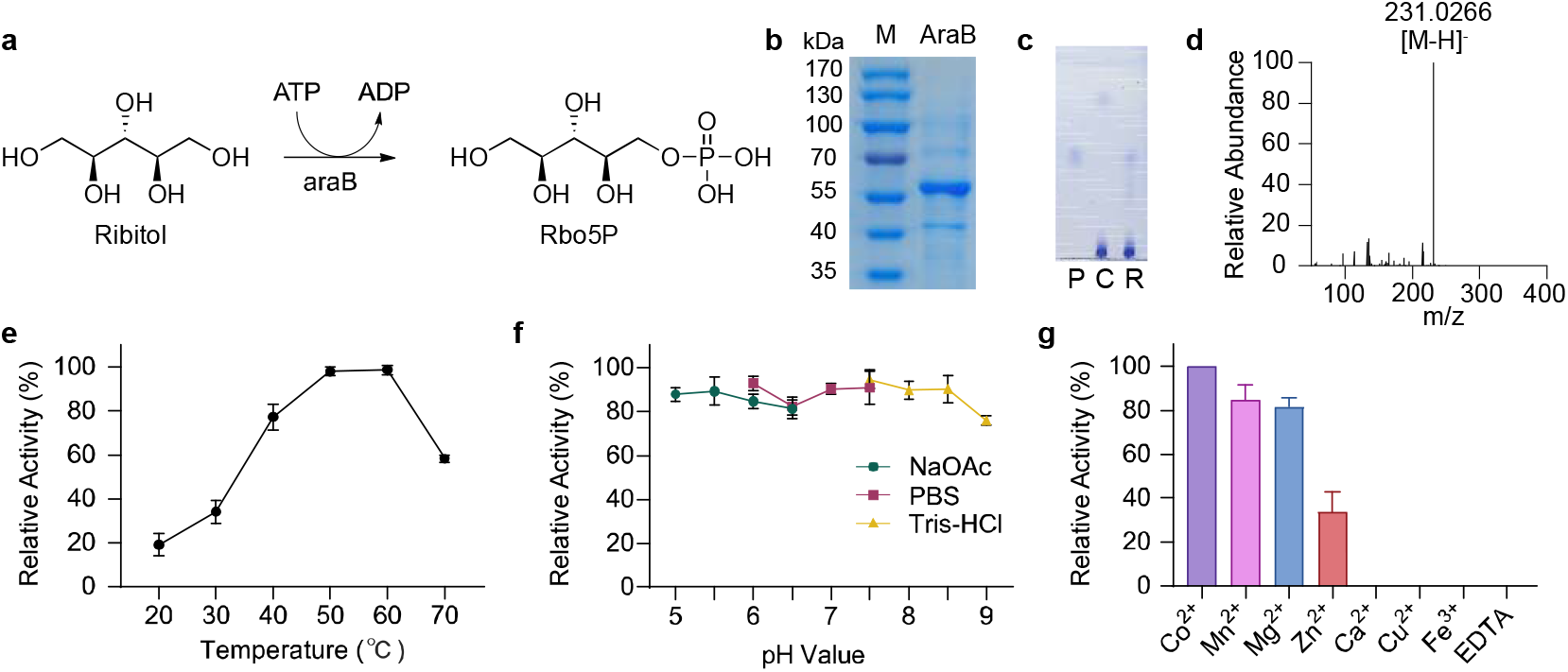
Phosphorylation of ribitol by AraB. (a) AraB catalyses the formation of Rbo5P from ribitol and ATP. (b) SDS-PAGE analysis of purified AraB, which migrates as a band at 62 kDa, consistent with its calculated molecular weight of 62.6 kDa. (c) TLC analysis of AraB activity. P. positive control, Rbo5P; C. control of reaction mixture without AraB; R. reaction mixture. (d) Mass spectrum of Rbo5P, the calculated [M–H]^−^ is *m/z* 231.0270, and an ion at *m/z* 231.0266 was observed in negative-ion ESI mode, confirming the formation of Rbo5P. (e) Relative activity of AraB from 20 to 70 °C (*n* = 3); the optimal temperature was 50 °C. (f) Relative activity of AraB from pH 5.0 to 9.0 (*n* = 3); the optimum was pH 7.5 (Tris-HCl). (g) Relative activity of AraB with different metal ions (*n* = 3); Co^2+^ was optimal.

The full-length *araB* gene was cloned from *E. coli* MG1655 and inserted into pET-28a with a C-terminal His-tag. The construct was expressed in *E. coli* BL21(DE3), and recombinant AraB was purified by Ni-NTA affinity chromatography, yielding 80.3 mg/L of protein with a molecular weight approximately 62.6 kDa (Fig. 3b). The activity of AraB toward ribitol was assayed in a reaction mixture containing 50 mM Tris-HCl (pH 7.0), 10 mM ribitol, 15 mM ATP and 20 mM MgCl_2_ at 37 °C. After 30 min, ribitol was fully consumed (Fig. 3c). Mass spectrometry analysis revealed a new product with an [M–H]^−^ ion at m/z 231.0266 (Fig. 3d), consistent with the mass of Rbo5P.

The optimal temperature, pH, and metal ion of AraB were then determined. The enzyme showed maximal activity at 50– 60 °C and across a broad pH range of 5–9 (Fig. 3e, f). A divalent metal ion was also required for its activity, as EDTA treatment completely abolished enzymatic activity. Among the metal ions tested, Co^2+^ supported the highest activity (Fig. 3g). We next examined the substrate promiscuity of AraB using a panel of monosaccharides and sugar alcohols (Table 1). Besides ribitol, AraB accepted D-ribulose, D-arabitol, L-arabitol, and erythritol as substrates, but showed no activity toward D-mannitol, D-ribose, or xylitol, as evidenced by TLC analysis (Fig. S1).

**Table 1.**
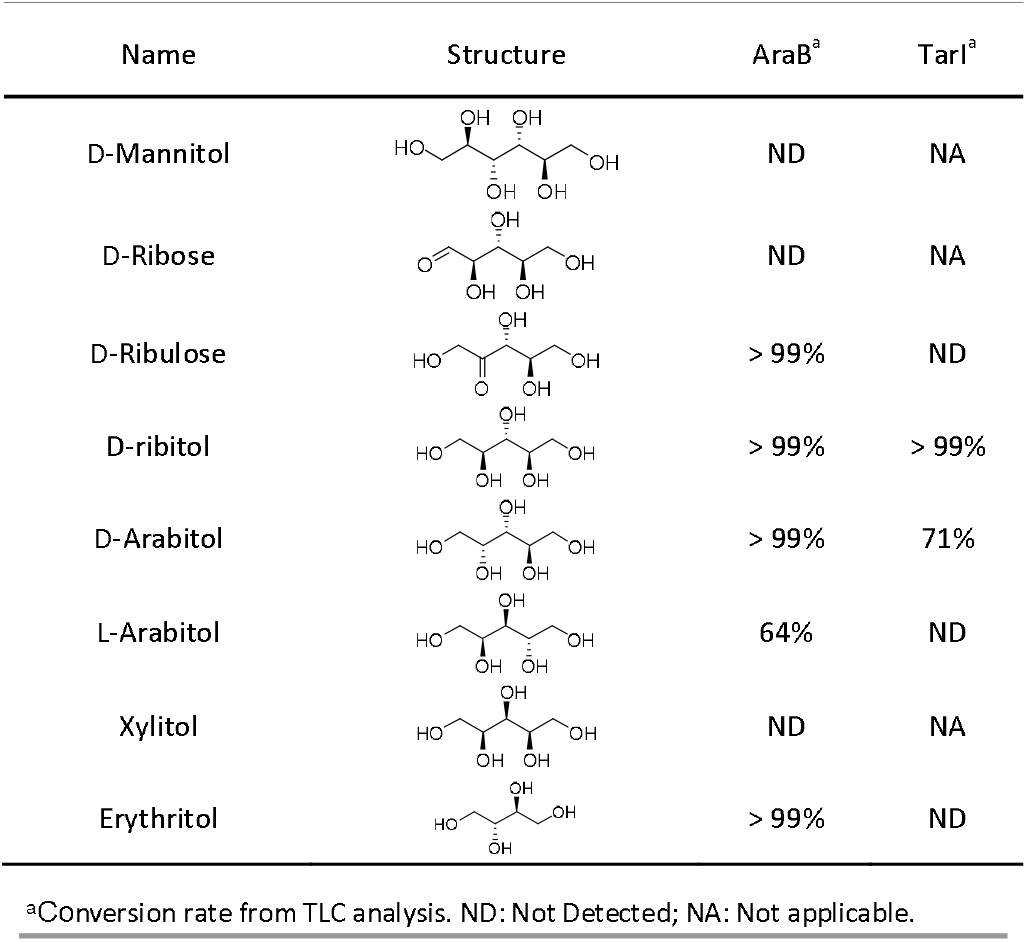
Activity assay of AraB and TarI

| Name | Structure | AraB<sup>a</sup> | TarI<sup>a</sup> |
| --- | --- | --- | --- |
| D-Mannitol |  | ND | NA |
| D-Ribose |  | ND | NA |
| D-Ribulose |  | > 99% | ND |
| D-ribitol |  | > 99% | > 99% |
| D-Arabitol |  | > 99% | 71% |
| L-Arabitol |  | 64% | ND |
| Xylitol |  | ND | NA |
| Erythritol |  | > 99% | ND |

<sup>a</sup>Conversion rate from TLC analysis. ND: Not Detected; NA: Not applicable.

We then scaled up the reaction for the synthesis of Rbo5P, using 200 mM Tris-HCl (pH 7.0), 200 mM ribitol, 300 mM ATP, 200 mM MgCl_2_ and 1 mg mL^−1^ AraB at 50 °C. Ribitol was fully converted to Rbo5P within 30 min. After purification by ion-exchange and Bio-Gel P-2 chromatography, 293 mg of Rbo5P was obtained in 90.6% isolated yield. Additionally, L-arabitol 5-phosphate (8 mg, 85%), D-arabitol 5-phosphate (19 mg, 46%), and erythritol 4-phosphate (22 mg, 86.3%) were also prepared. The structures of all synthesized products were confirmed by NMR spectroscopy, demonstrating the feasibility of this synthetic approach.

### Enzymatic synthesis of CDP-Rbo by TarI

CDP-Rbo pyrophosphorylase, also known as Rbo5P cytidylyltransferase, has been identified in many species (Fig. 4a), such as TarI in *S. pneumoniae*,^20^ *S. aureus*,^22^ *Bacillus subtilis*,^32^ Bcs1 in *H. influenzae*, ^33^ and ISPD in *H. sapiens*.^1, 4, 34^

**Fig. 4.**
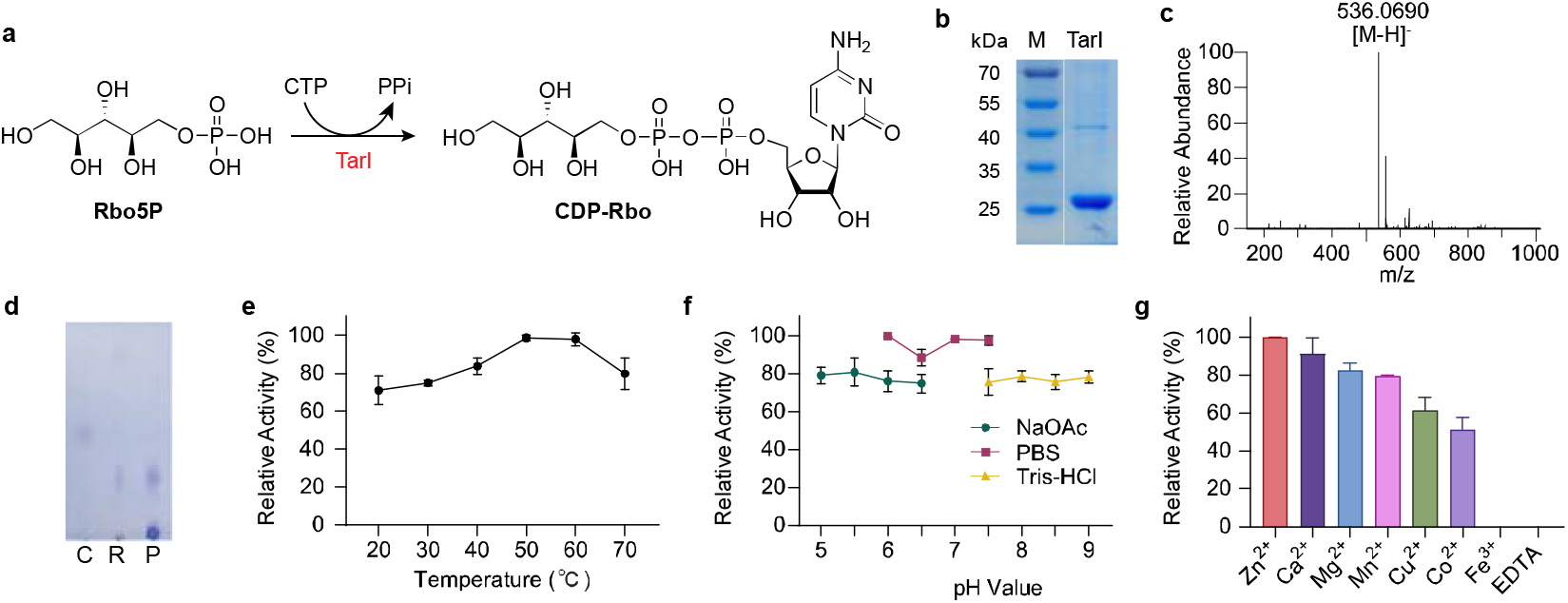
Synthesis of CDP-Rbo. (a) TarI-catalysed formation of CDP-Rbo from Rbo5P and CTP. (b) SDS-PAGE analysis of purified TarI, which migrates as a band at 27 kDa, consistent with its calculated molecular weight of 27.3 kDa. (c) ESI-MS of the product, the calculated [M–H]^−^ of CDP-Rbo is *m/z* 536.0682, and an ion at *m/z* 536.0690 was observed in negative-ion mode, confirming the formation of CDP-Rbo. (d) TLC analysis of TarI activity. P. positive control, CDP-Rbo; C. control of reaction mixture without TarI; R. reaction mixture. (e) Effect of temperature on TarI activity (*n* = 3); the optimum was 50-60 °C. (f) Effect of pH on TarI activity in sodium acetate, PBS and Tris-HCl buffers (*n* = 3); the optimum was PBS, pH 6.0. (g) Effect of metal ions on TarI activity (*n* = 3).

CDP-Rbo pyrophosphorylase (TarI) from *S. pneumoniae* was synthesized and inserted into pET28a with a C-terminal His-tag.^20^ The recombinant protein was expressed in *E. coli* BL21(DE3) at a level of 148.7 mg/L with a molecular weight approximately 27.3 kDa (Fig. 4b). The activity of TarI was assayed in a reaction mixture with 50 mM Tris-HCl (pH 7.0), 10 mM Rbo5P, 15 mM CTP, 20 mM MgCl_2_. The reaction was incubated at 37 °C and monitored by TLC analysis. After 30 min, the spot of Rbo5P disappeared and a new spot of CDP-Rbo was detected by TLC (Fig. 4d), which was confirmed by MS as an [M-H]^-^ ion at *m/z* 536.0690 (Fig. 4c).

The optimal temperature, pH, and metal-ion requirement of TarI were then measured. The enzyme was most active at 50–60 °C, with maximal activity observed in PBS buffer at pH 6.0–7.5 (Fig. 4e, f). A divalent metal ion was also essential for its activity, as EDTA treatment completely abolished it; among the metal ions tested, Zn^2+^ supported the highest activity (Fig. 4g). In contrast to the broad scope of AraB, TarI accepted only D-arabinitol 5-phosphate as substrate in addition to Rbo5P (Fig. S2).

CDP-Rbo was then prepared on a preparative scale in a reaction containing 200 mM PBS (pH 6.0), 200 mM Rbo5P, 300 mM CTP, 200 mM MgCl_2_ and 1 mg mL^−1^ TarI at 50 °C. Rbo5P was fully converted to CDP-Rbo after an overnight reaction. After purification by ion-exchange and Bio-Gel P-2 chromatography, 159 mg of CDP-Rbo was obtained in 90% isolated yield and characterized by NMR. Additionally, CDP-D-arabitol (58 mg, 53%) was prepared and characterized.

### Design and Evaluation of the bifunctional ribitol-dependent CDP-Rbo synthase AraB-TarI

Bifunctional enzymes that integrate two catalytic domains are widespread in nature, such as L-fucokinase/GDP-L-fucose pyrophosphorylase (FKP), which produces GDP-fucose from fucose.^35^ Such enzymes channel the phosphorylated product directly into the pyrophosphorylation step, without releasing it into solution. Inspired by these natural architectures, fusion enzymes have been engineered to facilitate sugar-nucleotide biosynthesis.^36-38^ For example, fusion of an N-acetylhexosamine 1-kinase with a truncated uridyltransferase allows the one-pot conversion of GlcNAc into UDP-GlcNAc.^36^ Such fused enzymes streamline multi-step reactions and provide a useful platform for preparing sugar-nucleotide analogues. Nevertheless, to date, no single fusion protein or naturally occurring bifunctional enzyme has been described for the direct conversion of ribitol to CDP-Rbo. Given that AraB and TarI share overlapping optimal temperatures and pH ranges, and both require a divalent metal ion for activity, we fused AraB with TarI to achieve a one-step synthesis of CDP-Rbo from ribitol.

AraB was fused to TarI through a flexible glycine/serine linker (Fig. 5a). Given the relatively large size of the fusion protein (89 kDa), we anticipated that soluble expression might be limiting and therefore evaluated two strategies in parallel: (i) a tag-free fusion, and (ii) an N-terminal maltose-binding protein (MBP) tagged fusion intended to enhance solubility (Fig. S3). For the MBP construct, four linkers (GGD, AGE, GGGRG and GSGGGSGHM) were compared to assess the effect of linker length and composition on activity.

**Fig. 5.**
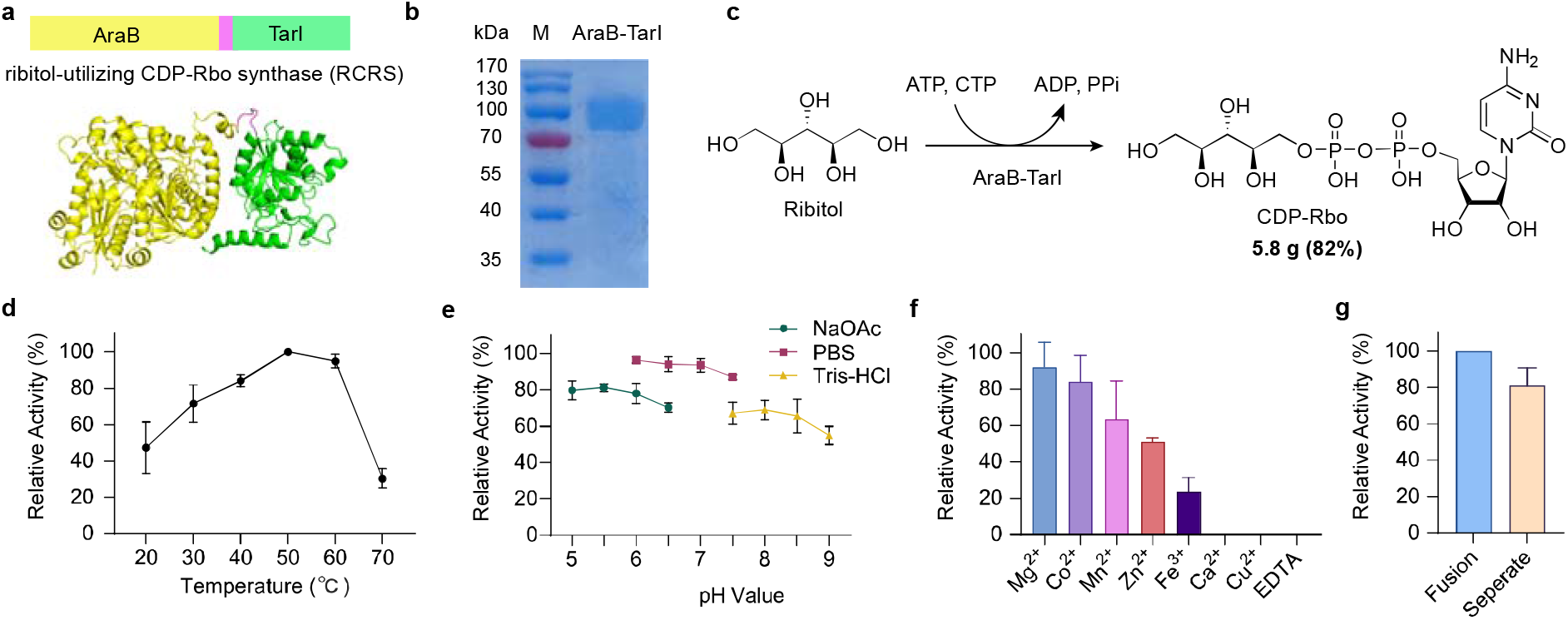
Bifunctional ribitol-dependent CDP-Rbo synthase AraB-TarI. (a) Design of AraB-TarI. Yellow color: AraB, green color: TarI, pink color: glycine and serine linker. The structure of AraB-TarI was predicted by AlphaFold 3.0. (b) SDS-PAGE analysis of purified AraB-TarI, which migrates as a band at 90 kDa, consistent with its calculated molecular weight of 89 kDa. (c) One-step reaction catalysed by AraB-TarI. (d) Effect of temperature on AraB-TarI activity (*n* = 3); the optimum was 50 °C. (e) Effect of pH on AraB-TarI activity in sodium acetate, PBS and Tris-HCl buffers (*n* = 3); the optimum was PBS, pH 6.0. (f) Effect of metal ions on AraB-TarI activity (*n* = 3). Mg^2+^ was optimal. (g) Activity comparison of AraB-TarI fusion enzyme with separate AraB and TarI. The reaction was conducted with equal enzyme molar amount.

After expression and purification, the expression level and purity of the target proteins were assessed. Contrary to our expectation, the tag-free fusion gave both the highest soluble expression (62.1 mg/L) and the highest purity, whereas the MBP-tagged constructs expressed at only 21–53 mg/L and were less pure by SDS-PAGE analysis (Fig. 5b, S4). The reduced expression observed for the MBP-tagged proteins (>130 kDa) is consistent with the general trend that soluble expression yields in *E. coli* tend to decrease with increasing protein size, as the MBP-tagged constructs (>130 kDa) are considerably larger than the tag-free fusion (89 kDa).^39^

The activity of the fusion enzymes was tested in a reaction mixture containing 50 mM Tris-HCl (pH 7.0), 10 mM ribitol, 20 mM MgCl_2_, 15 mM ATP and 15 mM CTP (Fig. 5c). The same amount of each fusion enzyme was added, and the mixture was incubated at 37 °C for 30 min. TLC analysis showed that ribitol was fully consumed by all fusion enzymes, although some Rbo5P remained unconverted to CDP-Rbo (Fig. S5). The MBP-tagged enzymes with different linkers gave similar conversion rates, indicating that linker length and composition had little effect on enzyme activity. As the tag-free fusion enzyme showed the highest expression and activity, it was named AraB-TarI and used for all subsequent studies.

The optimal temperature of AraB-TarI was 50 °C (Fig. 5d). AraB-TarI was most active in PBS buffer (pH 6.0), with Mg^2+^ as the optimal metal ion (Fig. 5e, f). We next compared the catalytic efficiency of the fused AraB-TarI with that of the individual AraB and TarI enzymes. At an equimolar enzyme concentration, the fused AraB-TarI achieved a higher conversion of ribitol to CDP-Rbo than an equivalent mixture of separate AraB and TarI, suggesting that fusion enzyme facilitates the transfer of Rbo5P from the catalytic domain of AraB to that of TarI (Fig. 5g). A gram-scale reaction was then performed using 2.0 g of ribitol as the starting material. The pH was adjusted to 6.0, and the mixture was incubated at 50 °C overnight. After purification by ion precipitation, 5.8 g CDP-Rbo was obtained in 82% isolated yield with >95% purity.^40^

### Nucleotide regeneration for the synthesis of CDP-Rbo

In the method described above, ATP and CTP are both consumed stoichiometrically (Fig. 6a), giving an atom economy of only 47%, in which the spent nucleotides (ADP and PPi) account for the majority of the lost mass. To improve the atom economy and the overall sustainability of the process, we therefore designed a cofactor-regeneration strategy.

**Fig. 6.**
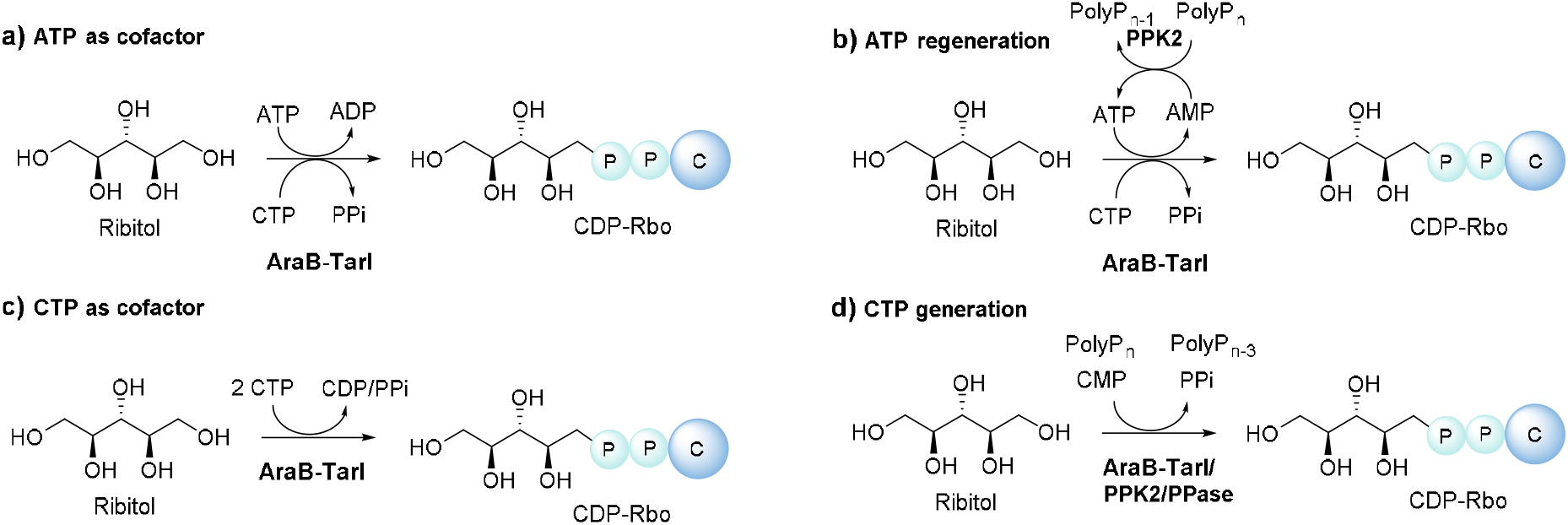
Strategies for the enzymatic synthesis of CDP-Rbo with nucleotide regeneration. All reactions are catalysed by the bifunctional AraB-TarI. (a) Synthesis with stoichiometric ATP and CTP as co-substrates (atom economy 47%). (b) Synthesis with catalytic ATP (or AMP), regenerated from ADP by polyphosphate kinase (PPK2) at the expense of inexpensive inorganic polyphosphate (polyPn); inorganic pyrophosphatase (PPase) hydrolyses the PPi released by the TarI domain to drive the reaction forward (atom economy 75%). (c) Synthesis using CTP as the sole nucleotide, in which CTP serves both as the phosphoryl donor for the AraB domain, releasing CDP, and as the cytidylyl donor for the TarI domain. (d) Synthesis from low-cost CMP, with CTP regenerated *in situ* by PPK2 and polyPn.

Polyphosphate kinase (PPK2) from *Erysipelotrichaceae* was introduced to regenerate ATP from ADP at the expense of inexpensive inorganic polyphosphate (polyP_n_) (Fig. 6b).^41^ The regeneration system was examined with ATP-to-ribitol ratios ranging from 1:5 to 1:100. After overnight reaction, ribitol was completely consumed in all reactions tested even at a ratio of 1:100 (Fig. S6a). Further lowering the ATP loading to 1:500 still afforded full conversion of ribitol to Rbo5P (Fig. S6b). Each ATP molecule was therefore recycled up to 500 times, corresponding to a total turnover number (TTN) of 500 (moles of Rbo5P per mole of ATP), compared with 0.67 for the non-regenerating route employing 1.5 equiv of ATP.

As PPK2 can phosphorylate AMP to ATP, the adenine nucleotide could be supplied as AMP rather than ATP. Thus, a regeneration reaction was tested with AMP-to-ribitol ratio of 1:100. Interestingly, AMP also gave a complete conversion of ribitol to Rbo5P (Fig. S6c). With this nucleotide regeneration strategy, the atom economy of the reaction increased from 47% to 75%. The regeneration also lowers the reagent cost, as polyP ($0.1/mmol) is two orders of magnitude cheaper than ATP ($11/mmol). Supplying the nucleotide as AMP ($2/mmol) reduces this cost a further five-fold.

Notably, although ribitol was completely converted to Rbo5P, the subsequent conversion of Rbo5P to CDP-Rbo was lower than expected. Because the TarI-catalysed cytidylylation is a reversible reaction, we reasoned that the accumulation of inorganic pyrophosphate (PPi) released during CDP-Rbo formation shifted the equilibrium and inhibited TarI. Accordingly, inorganic pyrophosphatase (PPase) from *Pasteurella multocida* was added to hydrolyze PPi, which drove the reaction forward with no detectable Rbo5P (Fig. S6d).^42^

Sugar kinases often show relaxed specificity towards the phosphoryl donor, so we tested whether the AraB domain would accept CTP instead of ATP. If so, a single cytidine nucleotide would serve both the kinase and the pyrophosphorylase steps (Fig. 6c). A reaction containing only ribitol and CTP was therefore performed. CDP-Rbo was detected by TLC within 0.5 h and accumulated over 2.5 h (Fig. S7), confirming that CTP is accepted by the AraB domain as a phosphoryl donor for the generation of Rbo5P from ribitol.

As CTP ($146/mmol) is more expensive than CMP ($7/mmol), we next combined CMP with PPK2 and polyPn to prepare CDP-Rbo instead of using expensive CTP (Fig. 6d). A reaction mixture containing 10 mM ribitol, 10 mM CMP, 20 mM MgCl_2_, 100 mM polyP_n_, PPK2, PPase, and AraB-TarI was incubated at 50 °C for 10 h. Ribitol was fully converted to Rbo5P, which was subsequently transformed into CDP-Rbo with conversion rate of 80.4% (Fig. S8). This strategy allows CDP-Rbo to be produced efficiently and at low cost, and should facilitate the large-scale preparation of CDP-Rbo and its analogues. Finally, it is worth noting that if this regeneration system were coupled to a Rbo5P glycosyltransferase, the CMP released upon glycosyl transfer would be re-phosphorylated to CTP, so that only a catalytic amount of CMP would be required. Under such conditions ribitol and polyP_n_ would be the only stoichiometric inputs, and both the atom economy and the TTN would increase further.

## Conclusions

In this work, we developed a practical strategy to produce CDP-Rbo from low-cost and readily available ribitol. An L-ribulokinase (AraB) and a CDP-Rbo pyrophosphorylase (TarI) were cloned and characterized to synthesize CDP-Rbo from ribitol via Rbo5P. Fusing the two enzymes gave a bifunctional CDP-Rbo synthase, AraB-TarI, which converts ribitol directly into CDP-Rbo in a single step. Using this synthase, CDP-Rbo was prepared on a 5.8 g scale in 82% isolated yield. To improve the sustainability, PPK2 and inexpensive inorganic polyphosphate were introduced to regenerate the nucleotides. The total turnover number of ATP could reach 500 and the atom economy of the production of CDP-Rbo was raised from 47% to 75%. Replacing CTP with the much cheaper CMP, which is phosphorylated *in situ* by PPK2 and polyphosphate, further reduced the reagent cost. The route thus provides efficient and economical access to CDP-Rbo, and a reliable material supply both for developing therapeutics against ISPD-deficient muscular dystrophy and for studying the Rbo5P-containing glycans of bacterial pathogens. Coupling this regeneration system to a Rbo5P glycosyltransferase would further increase the atom economy and reduce the cost, giving access to Rbo5P-containing glycans from ribitol and polyphosphate alone.

## Experimental

### Cloning and expression of AraB and TarI

The *araB* gene (UniProt P08204) was amplified from *E. coli* MG1655 and cloned into pET-28a using primers GGACAGCAAATGGGTCGCATGGCGATTGCAATTGGC and TTTGTTAGCAGCCGGATCTTATAGAGTCGCAACGGC. The TarI gene (UniProt Q8DPI2) was synthesized by Gene Universal and cloned into pET-28a. The resulting constructs were transformed into *E. coli* BL21(DE3) for expression.

The recombinant strains were grown in LB medium (10 g/L tryptone, 5 g/L yeast extract, 10 g/L sodium chloride) containing 100 μg/mL antibiotic at 37 °C overnight. The culture was inoculated into 1 L of LB medium and grown until the OD_600_ reached about 0.7. IPTG was then added to final concentration of 0.4 mM to induce protein expression, and the culture was incubated at 16 °C for about 20 h. Cells were harvested by centrifugation at 8000 rpm for 10 min, resuspended in PBS, and lysed by sonication. Cell debris was removed by centrifugation at 11 000 rpm for 30 min at 4 °C.

The supernatant was loaded onto a Ni-NTA column. Unbound proteins were removed with washing buffer (20 mM PBS, pH 7.5, 20 mM imidazole), and the target protein was eluted with elution buffer (20 mM PBS, pH 7.5, 300 mM imidazole). Protein purity was assessed by SDS-PAGE. Purified enzymes were stored in 20% (v/v) glycerol at −20 °C until use.

### Activity assay of AraB

The activity of AraB was assayed in a mixture containing 50 mM Tris-HCl (pH 7.0), 10 mM ribitol, 15 mM ATP, 20 mM MgCl_2_ and 50 μg/mL AraB. The mixture was incubated at 37 °C, monitored by TLC and confirmed by MS.

The optimal temperature was determined at 20, 30, 40, 50, 60 and 70 °C. Each reaction (20 μL) contained 10 mM ribitol, 15 mM ATP, 20 mM MgCl_2_ and 0.2 μg AraB at pH 7.0. The reaction was incubated at the corresponding temperature for 20 min. A negative control was prepared by replacing AraB with water.

The optimal pH was determined at pH 5.0 to 9.0 using 50 mM buffers: sodium acetate (pH 5.0, 5.5, 6.0, 6.5), PBS (pH 6.0, 6.5, 7.0, 7.5) and Tris-HCl (pH 7.5, 8.0, 8.5, 9.0). Each reaction (20 μL) contained 10 mM ribitol, 15 mM ATP, 20 mM MgCl_2_, the corresponding buffer, and 0.2 μg AraB. The reaction was incubated at 37 °C for 20 min. AraB was replaced with water in the negative control.

The optimal metal ion was determined using Na^+^, K^+^, Fe^3+^, Cu^2+^, Zn^2+^, Mn^2+^, Co^2+^, Mg^2+^ and EDTA. Each reaction (20 μL) contained 10 mM ribitol, 15 mM ATP, 20 mM of the corresponding metal ion or EDTA, and 0.2 μg AraB. The reaction was incubated at 37 °C for 20 min. AraB was replaced with water in the control. Reactions were stopped with ice-cold ethanol, and the product was quantified by TLC.

Substrate specificity was tested with D-mannitol, D-ribose, D-ribulose, D-arabitol, L-arabitol, xylitol and erythritol. Each reaction containing 50 mM Tris-HCl (pH 7.5), 20 mM of the corresponding substrate, 30 mM ATP, 40 mM MgCl_2_ and 50 μg/mL AraB was incubated at 37 °C, monitored by TLC and confirmed by MS and NMR. TLC plates were visualized with cerium ammonium molybdate (CAM) stain.

### Activity assay of TarI

The activity of TarI was assayed in a mixture containing 50 mM Tris-HCl (pH 7.0), 10 mM Rbo5P, 15 mM CTP, 20 mM MgCl_2_ and 50 μg/mL TarI. The mixture was incubated at 37 °C, monitored by TLC and confirmed by MS.

The optimal temperature was determined at 20, 30, 40, 50, 60 and 70 °C. Reactions (20 μL) containing 50 mM Tris-HCl (pH 7.0), 10 mM Rbo5P, 15 mM CTP, 20 mM MgCl_2_, and 0.4 μg TarI were incubated at the corresponding temperature for 20 min. A negative control was prepared by replacing TarI with water.

The optimal pH was determined at pH 5.0 to 9.0 using the 50 mM buffers described above. Each reaction (20 μL) contained 10 mM Rbo5P, 15 mM CTP, 20 mM MgCl_2_, the corresponding buffer and 0.4 μg TarI, and was incubated at 37 °C for 20 min. TarI was replaced with water in the negative control.

The optimal metal ion was determined using Fe^3+^, Cu^2+^, Zn^2+^, Mn^2+^, Co^2+^, Mg^2+^ and EDTA. Each reaction (20 μL) contained 10 mM Rbo5P, 15 mM CTP, 20 mM of the corresponding metal ion or EDTA, and 0.4 μg TarI, and was incubated at 37 °C for 20 min. TarI was replaced with water in the control. Reactions were stopped with ice-cold ethanol and quantified by TLC.

Substrate specificity was tested with D-ribulose 5-phosphate, D-arabitol 5-phosphate, L-arabitol 5-phosphate and erythritol 4-phosphate. Each reaction contained 50 mM PBS (pH 6.0), 20 mM of the corresponding substrate, 30 mM CTP, 40 mM MgCl_2_ and 50 μg/mL TarI, and was incubated at 37 °C, monitored by TLC and confirmed by MS and NMR.

### Construction of CDP-Rbo synthase

AraB and TarI were linked by a short peptide linker. Besides directed linked protein, an MBP tag was placed at the N-terminus. Four linkers were tested (GGD, AGE, GGGRG and GSGGGSGHM). The fusion sequences were cloned into the vector pMAL-c5X, and each construct carried a C-terminal His-tag for purification. Expression, purification, and activity assays of the fusion enzymes were performed as described above.

A gram-scale reaction was carried out using 2.0 g ribitol, 13.2 g ATP, 10.7 g CTP, 1.5 g MgCl_2_. The pH was adjusted to 6.0 and the reaction was incubated at 50 °C. After quench the reaction, ion precipitation method was used for purification^40^.

### Enzymatic synthesis of CDP-Rbo with nucleotide regeneration

To examine the ATP regeneration system, reactions were carried out at fixed concentrations of 40 mM ribitol, 50 mM CTP, 30 mM MgCl_2_, 500 mM polyP_n_ and 100 mM Tris-HCl (pH 7.5), with 5 mg mL^−1^ PPK2, 2 mg mL^−1^ AraB-TarI. ATP was added at ATP-to-ribitol molar ratios ranging from 1:5 to 1:500, corresponding to ATP concentrations of 8 mM (1:5), 4 mM (1:10), 2 mM (1:20), 1.33 mM (1:30), 1 mM (1:40), 0.8 mM (1:50), 0.67 mM (1:60), 0.57 mM (1:70), 0.5 mM (1:80), 0.44 mM (1:90), 0.4 mM (1:100), 0.2 mM (1:200), 0.13 mM (1:300), 0.1 mM (1:400) and 0.08 mM (1:500). Reactions were incubated at 50 °C overnight and analyzed by TLC.

To replace ATP with AMP, reactions were prepared as above (40 mM ribitol, 50 mM CTP, 30 mM MgCl_2_, 500 mM polyP_n_, 100 mM Tris-HCl pH 7.5, 5 mg mL^−1^ PPK2, 2 mg mL^−1^ AraB-TarI) but with AMP in place of ATP, added at 0.4 mM (AMP-to-ribitol 1:100) or 0.08 mM (1:500). Reactions were incubated at 50 °C overnight and analyzed by TLC.

To achieve complete conversion of the Rbo5P to CDP-Rbo, PPase was used in the reaction mixture containing 10 mM Rbo5P, 10 or 20 mM CMP, 20 mM MgCl_2_, 100 mM polyP_n_ and 100 mM Tris-HCl (pH 7.5), with 5 mg mL^−1^ PPK2, 2 mg mL^−1^ AraB-TarI and 1 mg mL^−1^ PPase. Reactions were incubated at 50 °C overnight and analysed by TLC.

For CTP as the sole nucleotide, reactions contained 10 mM ribitol, 15 mM CTP, 20 mM MgCl_2_ and 100 mM Tris-HCl (pH 7.5), with 2 mg mL^−1^ AraB-TarI. Reactions were incubated at 50 °C, and aliquots were withdrawn at 0.5 h and 2.5 h for analysis.

For the one-pot synthesis from ribitol and CMP, reactions contained 10 mM ribitol, 10 mM CMP, 20 mM MgCl_2_, 100 mM polyP□ and 100 mM Tris-HCl (pH 7.5), with 5 mg mL^−1^ PPK2, 2 mg mL^−1^ AraB-TarI and 1 mg mL^−1^ PPase. Reactions were incubated at 50 °C overnight and analyzed by HPLC.

## Author contributions

X. Zhang: Investigation, Methodology, Data curation, Validation, Writing – original draft; L. Pan: Data curation, Validation, Visualization; Y. Wang, M. Bi, X. Liu, and P. Ling: Writing – review & editing; C. Chen and S. Wang: Conceptualization, Supervision, Writing – review & editing.

## Data availability

The data supporting this article have been included as part of the Supplementary Information. Supplementary information: the sources of important compounds, experimental protocols, Fig. S1–S8, Tables S1-S2, NMR and MS spectra.

## Acknowledgements

This work was supported by the National Key Research and Development Program of China (2023YFF1103600), National Natural Science Foundation of China (22407077), Taishan Scholars Program (TSQN202312003), Guangdong Basic and Applied Basic Research Foundation (2023A1515110619), Natural Science Foundation of Shandong Province (ZR2025MS175, ZR2025QC1313), Basic Research Program of Jiangsu (BK20250410), the Qilu Young Scholar Fund of Shandong University, and the Intramural Joint Program Fund of State Key Laboratory of Microbial Technology (SKLMTIJP-2025-03). We thank Haiyan Sui and Jingyao Qu of the Core Facilities for Life and Environmental Sciences, State Key Laboratory of Microbial Technology of Shandong University for NMR and MS analysis.

