## Supplemental File for "Gram-scale one-pot enzymatic synthesis of CDP-ribitol by a designed bifunctional fusion enzyme"

**Supplementary Figures and Tables**

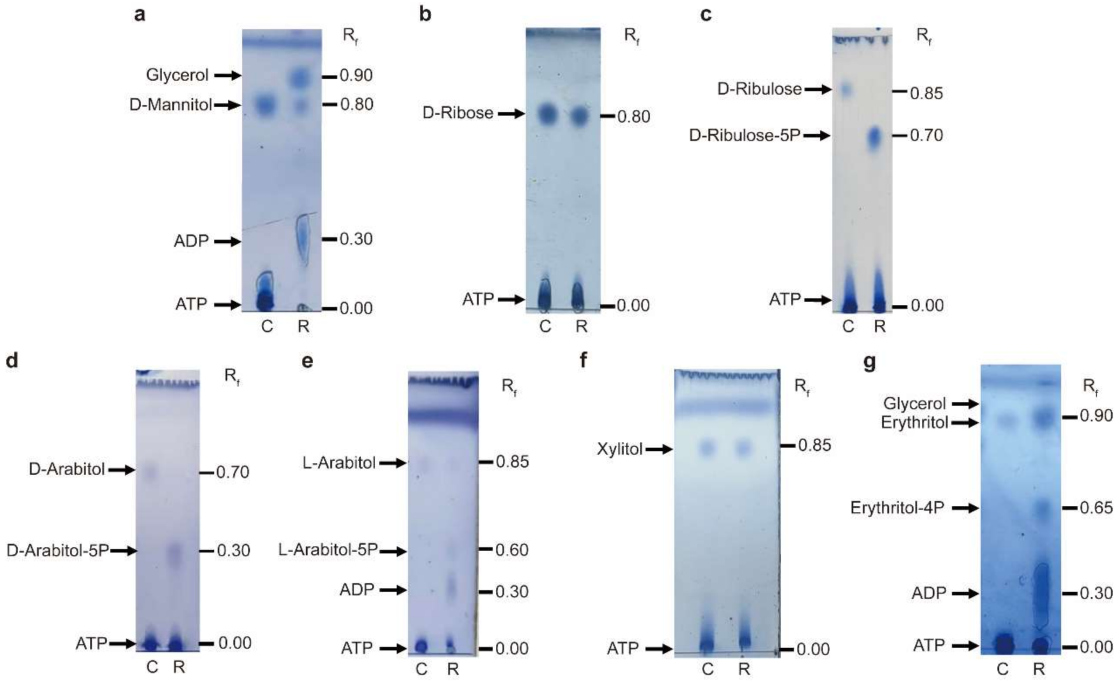


Fig. S1: AraB activity to ribitol analogs.

(a) When D-mannitol was used as the substrate, no product spot was observed by TLC. (b) When D-ribose was used as the substrate, no product spot was observed by TLC. (c) When D-ribulose was used as the substrate, a product spot corresponding to product D-ribulose-5P was detected by TLC at $R_f$ = 0.70. (d) When D-arabitol was used as the substrate, a product spot corresponding to product D-arabitol-5P was detected by TLC at $R_f$ = 0.30. (e) When L-arabitol was used as the substrate, a product spot corresponding to product L-arabitol-5P was detected by TLC at $R_f$ = 0.60. (f) When Xylitol was used as the substrate, no product spot was observed by TLC. (g) When erythritol was used as the substrate, a product spot corresponding to product erythritol-4P was detected by TLC at $R_f$ = 0.65.

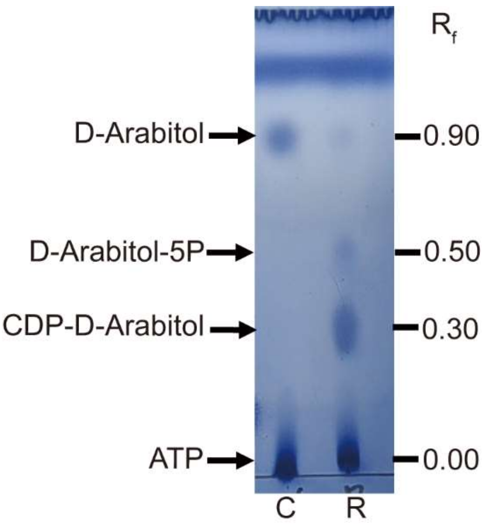


Fig. S2: TarI activity toward ribitol 5-phosphate analogs.

When D-arabitol was used as the substrate, D-arabitol-5P was detected by TLC at $R_f$ = 0.50 and CDP-D-arabitol was detected by TLC at $R_f$ = 0.30.

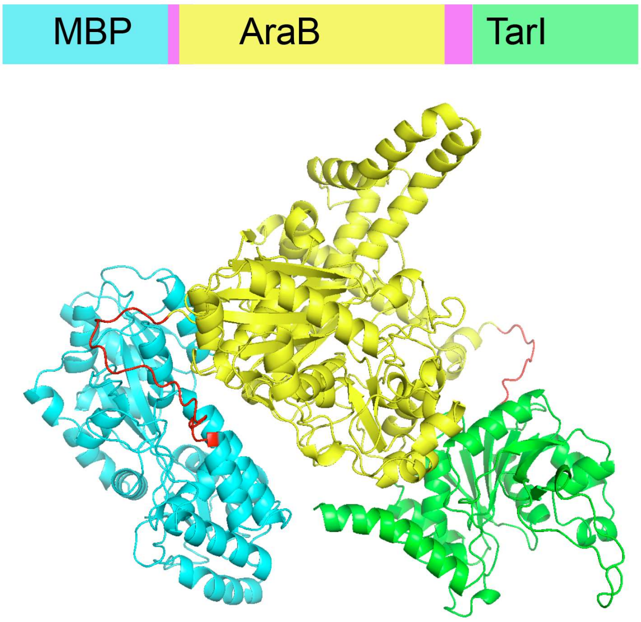


Fig. S3: AlphaFold 3 prediction of MBP-AraB-TarI fused enzyme.

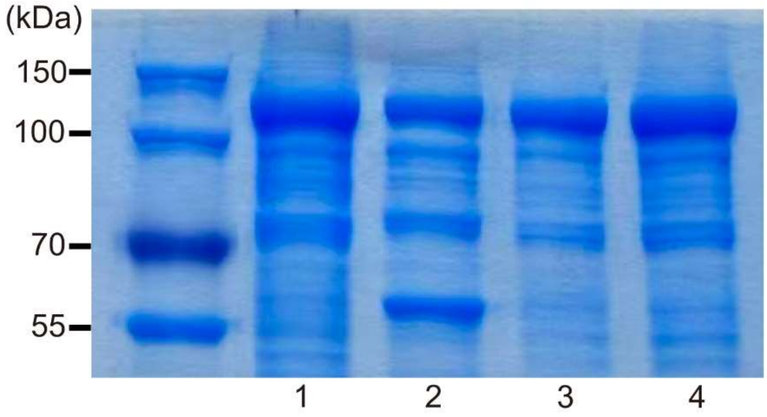


Fig. S4: SDS-PAGE analysis of MBP-AraB-TarI fused enzyme.

1 represents the SDS-PAGE of the enzyme with linker $NH_2$-GSGGGSGHM-COOH. 2 represents the SDS-PAGE of the enzyme with linker $NH_2$-GGGRG-COOH. 3 represents the SDS-PAGE of the enzyme with linker $NH_2$-GGD-COOH. 4 represents the reaction mixture of the enzyme with linker $NH_2$-AGE-COOH.

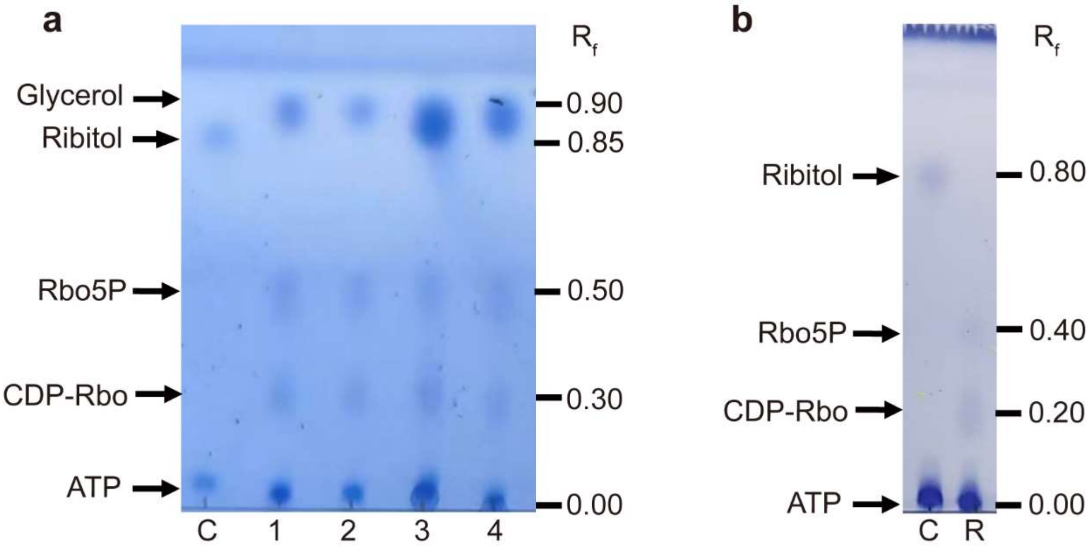


Fig. S5: Determination of the activity of fused bifunctional enzyme.

(a) TLC analysis of MBP-AraB-TarI fused enzyme. 1 represents the reaction mixture of the enzyme with linker $NH_2$-GGD-COOH. 2 represents the reaction mixture of the enzyme with linker $NH_2$-AGE-COOH. 3 represents the reaction mixture of the enzyme with linker $NH_2$-GGGRG-COOH. 4 represents the reaction mixture of the enzyme with linker $NH_2$-GSGGGSGHM-COOH. Rbo5P was detected by TLC at $R_f$ = 0.50 and CDP-Rbo at $R_f$ = 0.30. (b) TLC analysis of AraB-TarI fused enzyme. Rbo5P was detected by TLC at $R_f$ = 0.40 and CDP-Rbo at $R_f$ = 0.20.

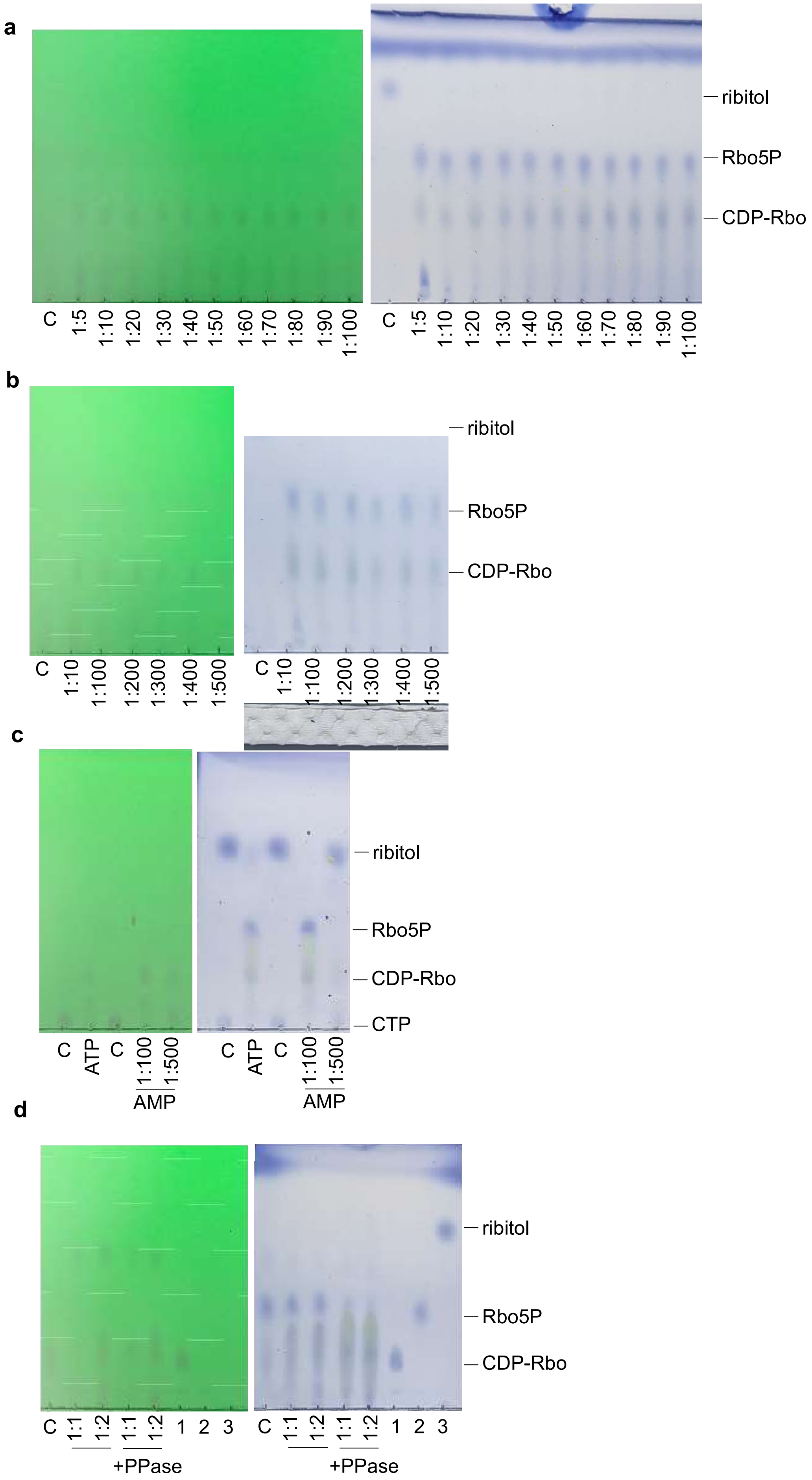
a
C
1:5
1:10
1:20
1:30
1:40
1:50
1:60
1:70
1:80
1:90
1:100
ribitol
Rbo5P
CDP-Rbo
b
1:200
1:300
1:400
1:500
c
ATP
1:100
1:500
AMP
CTP
d
1:1
1:2
1
2
3
+PPase

Fig. S6 Enzymatic synthesis of CDP-Rbo with ATP/AMP regeneration

(a) TLC analysis of reaction mixture with ATP to ribitol ratio from 1:5 to 1:100. (b) TLC analysis of reaction mixture with ATP to ribitol ratio from 1:10 to 1:500. (c) TLC analysis of ATP/AMP regeneration. ATP represent ATP to ribitol ratio as 1:500, AMP 1:100 and AMP 1:500 represent the AMP to ribitol ratio as 1:100 and 1:500, respectively. The ribitol to CTP ratio is 1:1. (d) TLC analysis of reaction mixture with/without PPase. The Rbo5P to CMP ratio is 1:1 and 1:2. The Rbo5P could be converted to CDP-Rbo with the employment of PPase.

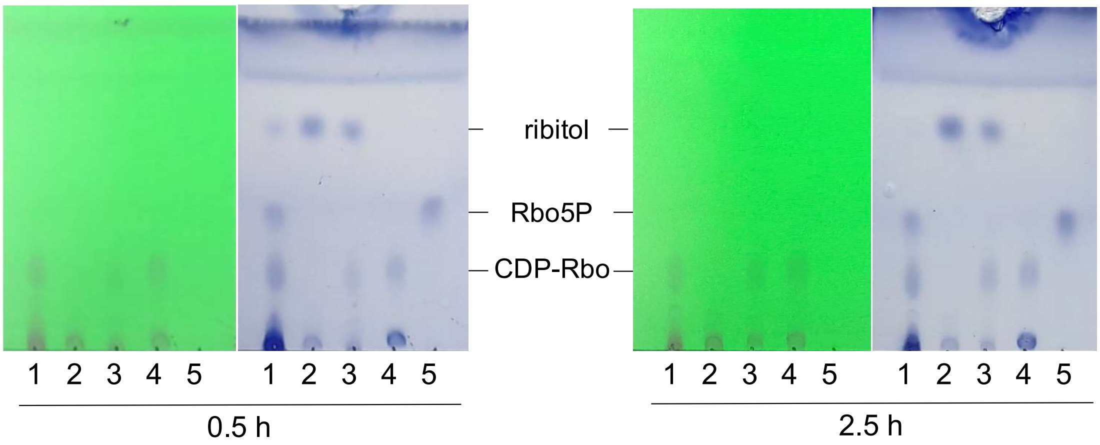


Fig. S7 TLC analysis result for enzymatic synthesis of CDP-Rbo with CTP.

Lane 1 reaction with ATP; Lane 2 negative control; Lane 3 reaction with CTP only; Lane 4 CDP-Rbo; Lane 5 Rbo5P

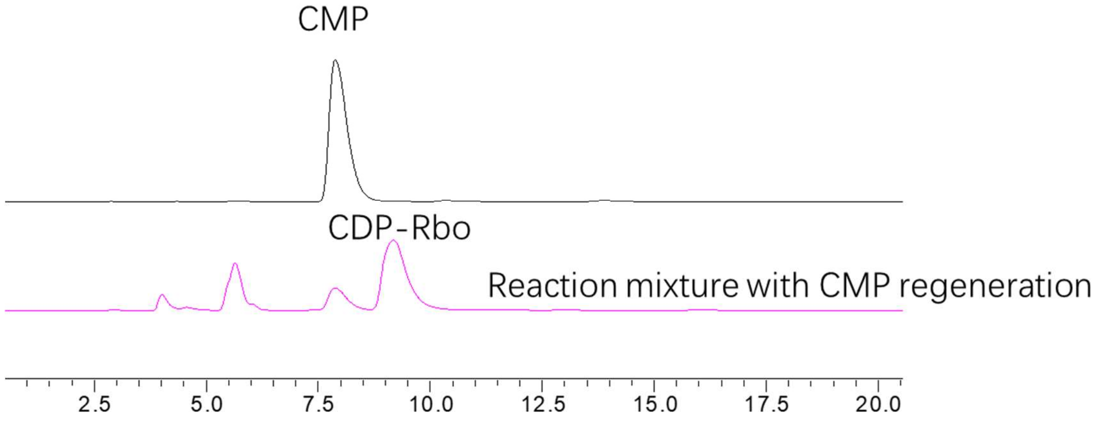


Fig. S8 HPLC analysis of the synthesis of CDP-Rbo from CMP regeneration.

Black: CMP; pink: reaction mixture with CMP regeneration. The conversion rate is 80.4%.

Table S1. Design and characterization of fused enzymes

| Fused protein | Constructs | Linker | Expression level (mg/L) | Conversion rate |
| --- | --- | --- | --- | --- |
| 1 | AraB-TarI | GSGGGHM | 62.1 | 66.9% |
| 2 | MBP-AraB-TarI | GSGGGSGHM | 46.5 | 43.9% |
| 3 | MBP-AraB-TarI | GGGRG | 21.6 | 57.1% |
| 4 | MBP-AraB-TarI | GGD | 53.2 | 50.3% |
| 5 | MBP-AraB-TarI | AGE | 30.1 | 46.1% |

Table S2. Prices of substrates and nucleotides from Fisher Scientific

| Abbreviation | Cat. No. | CAS NO. | List Price | MW (g/mol) | Price ($/mmol) |
|---|---|---|---|---|---|
| Ribitol | 167035000 | 488-81-3 | $1697/500 g | 152.15 | 0.52 |
| PolyP | 612175000 | 68915-31-1 | $96/500 g | 611.52 | 0.11 |
| AMP | J61643-14 | 4578-31-8 | $131/25 g | 391.19 | 2.05 |
| CMP | J63376-14 | 6757-06-8 | $475/25 g | 367.16 | 6.98 |
| ATP | J61125-14 | 34369-07-8 | $483/25 g | 551.15 | 10.65 |
| CTP | J62238-04 | 36051-68-0 | $555/2 g | 527.12 | 146.28 |

Maximum package size for nucleotide sodium salts.

**Supplementary methods**

### 1. Materials and Enzymes

All chemicals used in this study were of analytical grade. D-mannitol (CAS: 69-65-8), D-arabitol (CAS: 488-82-4) and D-ribulose (CAS: 488-84-6) were purchased from Shanghai Yuanye Bio-Technology Co., Ltd. D-ribose (CAS: 50-69-1) and Xylitol (CAS: 87-99-0) were obtained from Shanghai Dibai Biotechnology Co., Ltd. Erythritol (CAS: 149-32-6) was supplied by Shandong Sanyuan Biotechnology Co., Ltd. L-arabitol (CAS: 7643-75-6) was purchased from Aladdin Industrial Corporation, and ribitol (CAS: 488-81-3) was provided by Beijing psaitong Biotechnology Co., Ltd.

### 2. Experimental Procedures

#### 2.1 The cloning, expression and purification of AraB

The *araB* gene (UniProt P08204) was amplified from *E. coli* MG1655 using primers GGACAGCAAATGGGTCGCATGGCGATTGCAATTGGC and TTTGTTAGCAGCCGGATCTTATAGAGTCGCAACGGC. The gene was cloned into vector pET-28a with His tag at C-terminal. The plasmid of AraB was transformed into chemically competent cells of *E. coli* BL21(DE3) strain, and recombinant AraB was purified by Ni-NTA affinity chromatography.

#### 2.2 The cloning, expression and purification of AraB-TarI

The gene *araB* was cloned into TarI-containing pET-28a with C-terminal His-tag. Linker-GSGGGHM was introduced between the two genes, with primers CATATGATGATCTACGCCGGCATTCT, AATCGCCATGGTATATCTCCTTCTTAAAGTTAAACA, AAGAAGGAGATATACCATGGCGATTGCAATTG and CATATGGCCGCCGCCGCTGCCTAGAGTCGCAA applied. The recombinant AraB-TarI was purified by Ni-NTA affinity chromatography.

#### 2.3 The cloning, expression and purification of MBP-AraB-TarI

The *araB* gene was first cloned into the pMAL-c5X vector. Subsequently, gene TarI was inserted into the pMAL-c5X vector harboring gene *araB*. Linker GSGGGSGHM, GGGRG, GGD, AGE were designed between the two genes during the cloning of gene TarI. The plasmids of MBP-AraB-TarI with linker GSGGGSGHM, GGD and AGE were transformed into chemically competent cells of *E. coli* BL21(DE3) strain. The plasmids of MBP-AraB-TarI with linker

GGGRG was transformed into chemically competent cells of *E. coli* Rosetta(DE3) strain. The vector pMAL-c5X with MBP tag at N-terminal and His tag at C-terminal. One-step purification via MBP (Maltose-Binding Protein) affinity chromatography achieved a purified recombinant fusion protein.

2.4 Enzymatic synthesis

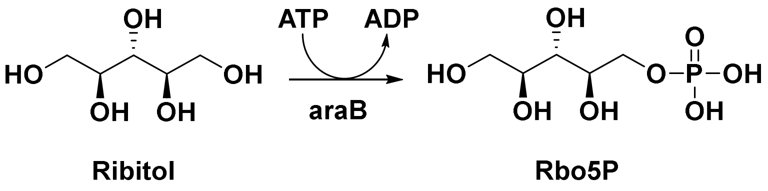


Synthesis of Rbo5P: Ribitol (10 mM, 1.0 equiv.), ATP (15 mM, 1.5 equiv.), $MgCl_2$ (20 mM), Tris-HCl buffer (50 mM, pH 7.0), AraB (50 μg/mL), 37 °C, 30 min. $^1$H NMR (600 MHz, $D_2O$) δ 3.99 (ddd, *J* = 11.5, 6.4, 3.1 Hz, 1H), 3.94 (dt, *J* = 11.5, 6.3 Hz, 1H), 3.88 (tq, *J* = 7.7, 4.5, 3.7 Hz, 2H), 3.84 – 3.78 (m, 1H), 3.76 (t, *J* = 6.3 Hz, 1H), 3.65 (dd, *J* = 11.8, 7.2 Hz, 1H); $^{13}$C NMR (151 MHz, $D_2O$) δ 72.11, 71.78, 71.23, 71.18, 65.23, 65.20, 62.33; HRMS (ESI) *m/z* calcd for $C_5H_{13}O_8P$ $[M-H]^-$ 231.0270, found 231.0265.

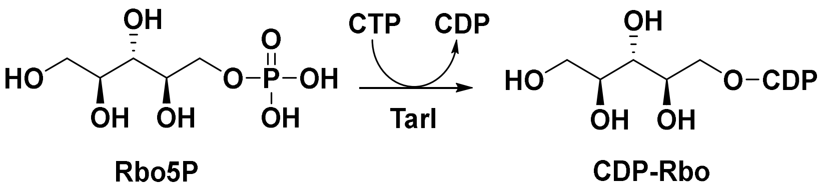


Synthesis of CDP-Rbo: Rbo5P (10 mM, 1.0 equiv.), CTP (15 mM, 1.5 equiv.), $MgCl_2$ (20 mM), Tris-HCl buffer (50 mM, pH 7.0), TarI (50 μg/mL), 37 °C, 30 min. $^1$H NMR (600 MHz, $D_2O$) δ 8.07 (dd, *J* = 7.8, 1.2 Hz, 1H), 6.21 (dd, *J* = 7.7, 1.2 Hz, 1H), 6.06 – 5.89 (m, 1H), 4.43 – 4.25 (m, 4H), 4.25 – 3.98 (m, 3H), 3.99 – 3.72 (m, 4H), 3.66 (ddd, *J* = 11.9, 7.2, 1.1 Hz, 1H); $^{13}$C NMR (151 MHz, $D_2O$) δ 165.10, 156.34, 141.83, 96.40, 89.24, 82.79, 82.73, 74.25, 72.05, 71.58, 70.86, 70.80, 69.24, 66.97, 66.93, 64.61, 64.57, 62.29; HRMS (ESI) *m/z* calcd for $C_{14}H_{25}N_3O_{15}P_2$ $[M-H]^-$ 536.0682, found 536.0687.

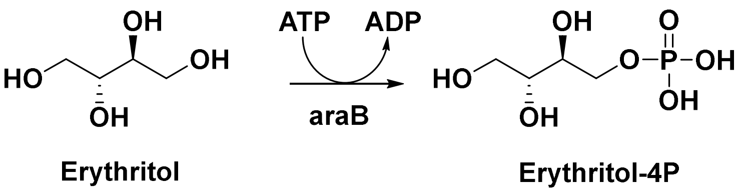


Synthesis of Erythritol-4P: Erythritol (10 mM, 1.0 equiv.), ATP (15 mM, 1.5 equiv.), $MgCl_2$ (20 mM), Tris-HCl buffer (50 mM, pH 7.0), AraB (50 μg/mL),

37 °C, 30 min. $^1$H NMR (600 MHz, $D_2O$) δ 3.96 (ttd, *J* = 11.4, 7.8, 7.2, 4.0 Hz, 2H), 3.82 (dd, *J* = 11.8, 2.8 Hz, 1H), 3.79 – 3.69 (m, 2H), 3.69 – 3.59 (m, 1H); $^{13}$C NMR (151 MHz, $D_2O$) δ 71.90, 71.31, 70.92, 70.87, 65.74, 65.71, 62.61, 62.58, 62.15; HRMS (ESI) *m/z* calcd for $C_4H_{11}O_7P$ $[M-H]^-$ 201.0146, found 201.0177.

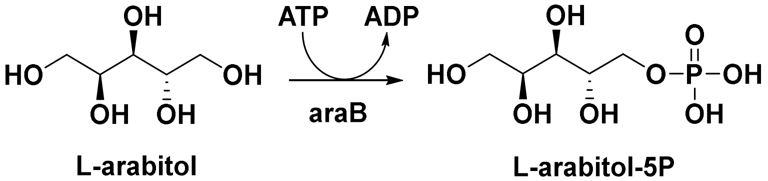


Synthesis of L-arabitol-5P: L-arabitol (10 mM, 1.0 equiv.), ATP (15 mM, 1.5 equiv.), $MgCl_2$ (20 mM), Tris-HCl buffer (50 mM, pH 7.0), AraB (50 µg/mL), 37 °C, 30 min. $^1$H NMR (600 MHz, $D_2O$) δ 4.07 – 3.93 (m, 3H), 3.81 (ddd, *J* = 8.8, 4.4, 2.8 Hz, 1H), 3.75 – 3.61 (m, 3H); $^{13}$C NMR (151 MHz, $D_2O$) δ 70.32, 70.28, 69.75, 65.31, 65.28, 63.16; HRMS (ESI) *m/z* calcd for $C_5H_{13}O_8P$ $[M-H]^-$ 231.0270, found 231.0276.

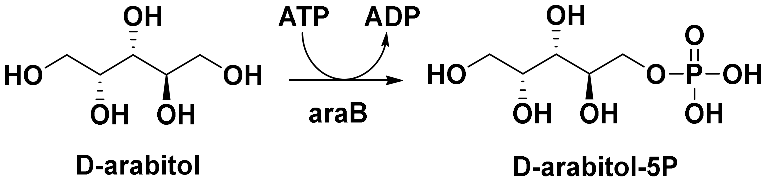


Synthesis of D-arabitol-5P: D-arabitol (10 mM, 1.0 equiv.), ATP (15 mM, 1.5 equiv.), $MgCl_2$ (20 mM), Tris-HCl buffer (50 mM, pH 7.0), AraB (50 µg/mL), 37 °C, 3 h. $^1$H NMR (600 MHz, $D_2O$) δ 3.96 (ddd, J = 11.3, 6.1, 2.7 Hz, 1H), 3.94 – 3.85 (m, 2H), 3.85 – 3.71 (m, 2H), 3.71 – 3.49 (m, 4H), $^{13}$C NMR (151 MHz, $D_2O$) δ 71.11, 71.07, 70.24, 70.17, 70.13, 69.74, 65.63, 65.60, 65.23, 65.19, 63.12, 62.20, HRMS (ESI) *m/z* calcd for $C_5H_{13}O_8P$ $[M-H]^-$ 231.0270, found 231.0260.

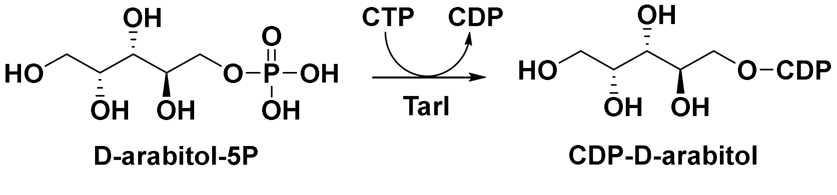


Synthesis of CDP-D-arabitol: D-arabitol-5P (10 mM, 1.0 equiv.), CTP (15 mM, 1.5 equiv.), $MgCl_2$ (20 mM), Tris-HCl buffer (50 mM, pH 7.0), TarI (50 µg/mL), 37 °C, 12 h. $^1$H NMR (600 MHz, $D_2O$) δ 8.10 (d, J = 7.8 Hz, 1H), 6.23 (d, J = 7.8 Hz, 1H), 5.98 (d, J = 4.1 Hz, 1H), 4.36 (dq, J = 9.2, 4.9 Hz, 2H), 4.31 (dtd, J = 6.9, 4.5, 2.4 Hz, 2H), 4.21 (ddt, J = 16.6, 8.0, 2.8 Hz, 2H), 4.12 (ddd, J =

11.1, 6.4, 4.9 Hz, 1H), 3.96 (ddd, J = 7.3, 5.3, 2.0 Hz, 1H), 3.88 (dtt, J = 7.6, 2.7, 1.4 Hz, 1H), 3.73 – 3.67 (m, 2H), 3.69 – 3.64 (m, 1H), $^{13}C$ NMR (151 MHz, $D_2O$) δ 162.74, 153.24, 142.74, 95.96, 89.36, 83.00, 82.94, 74.30, 70.16, 69.64, 69.59, 69.49, 69.17, 67.37, 67.33, 64.52, 64.48, 63.11, HRMS (ESI) *m/z* calcd for $C_{14}H_{25}N_3O_{15}P_2$ $[M-H]^-$ 536.0682, found 536.0683.

**MS and NMR spectra**

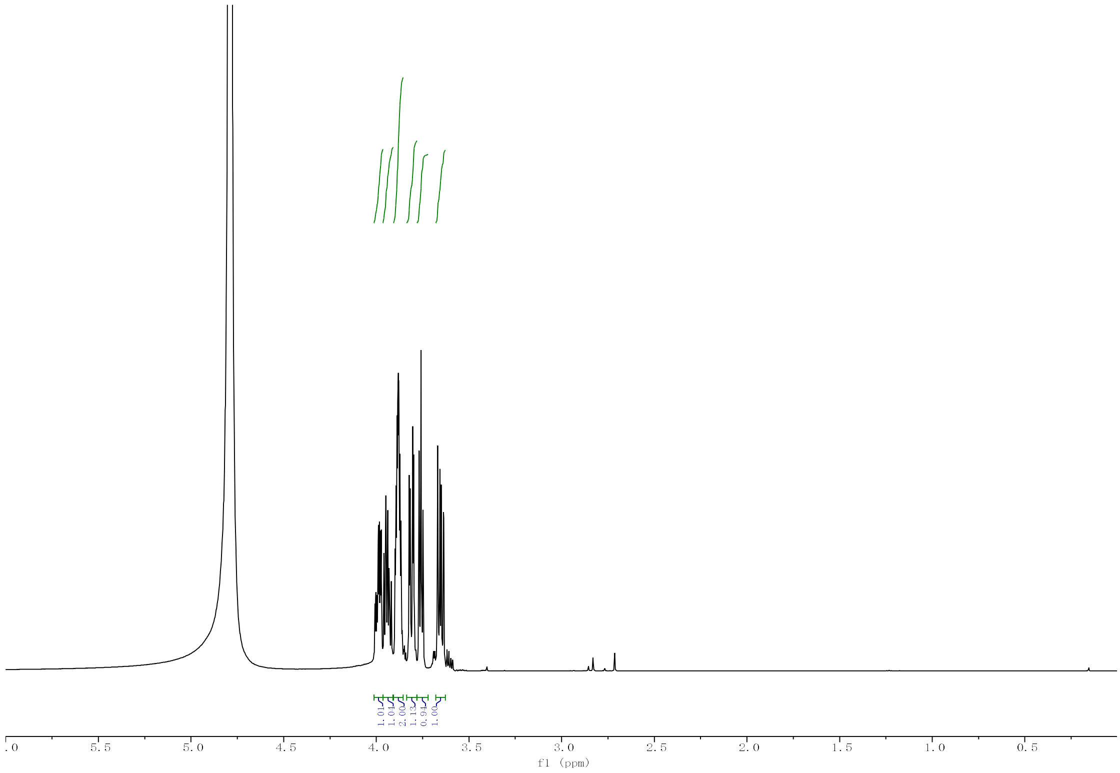


$^1H$ NMR of Rbo5P.

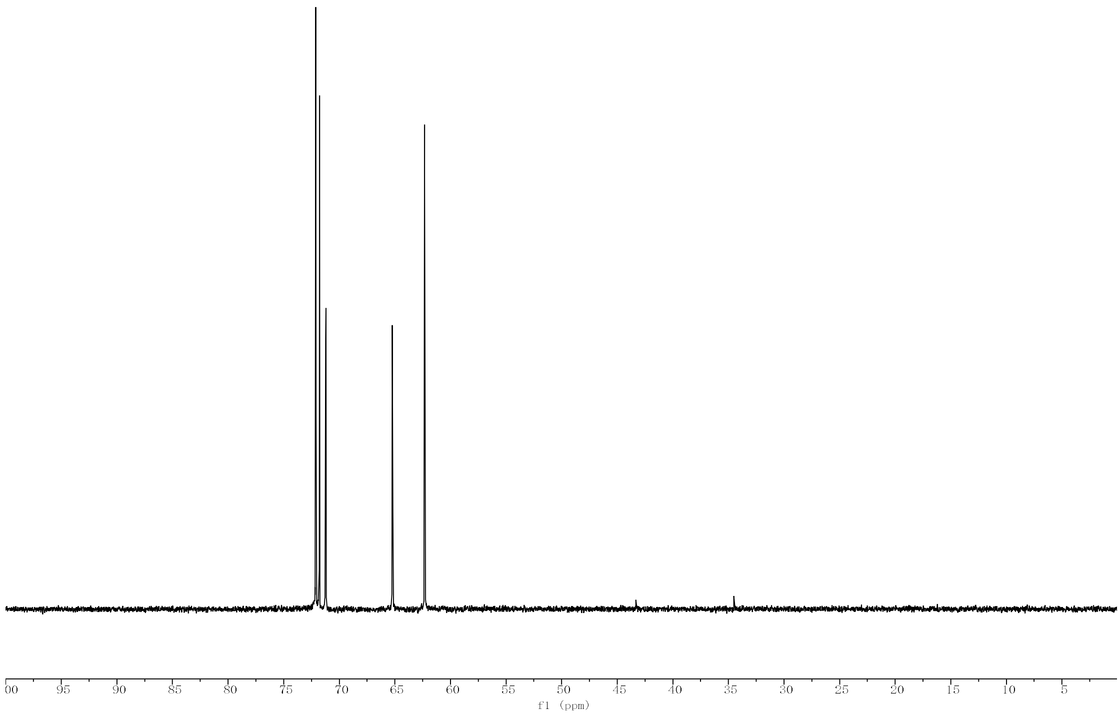


$^{13}C$ NMR of Rbo5P.

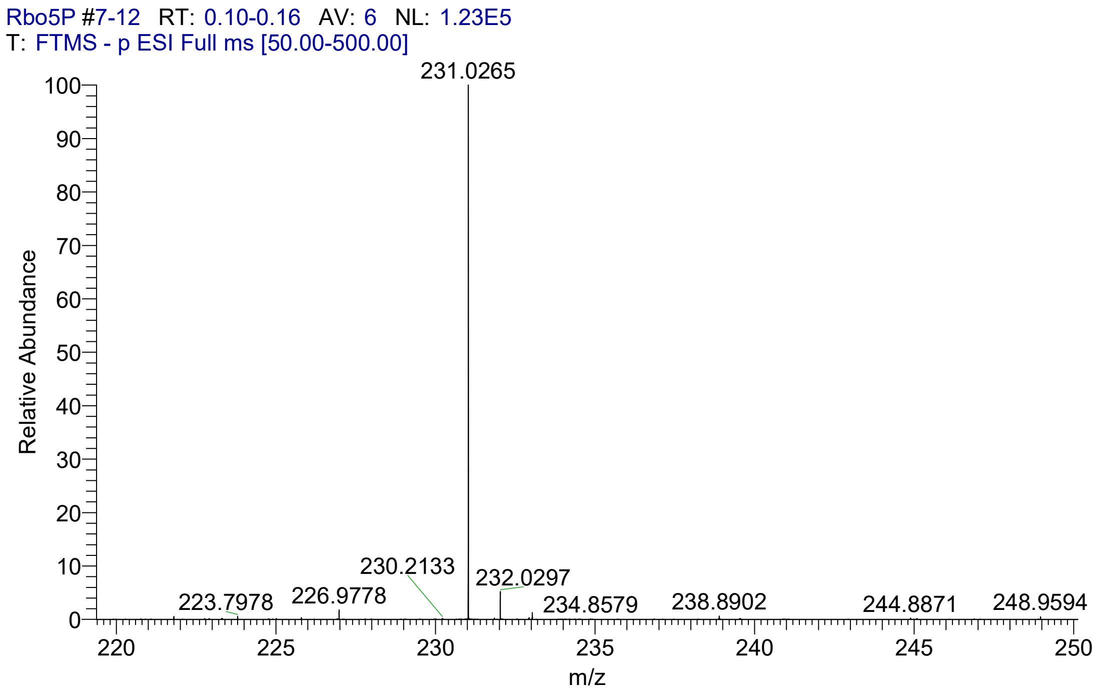
Rbo5P #7-12 RT: 0.10-0.16 AV: 6 NL: 1.23E5
T: FTMS - p ESI Full ms [50.00-500.00]
231.0265
230.2133
232.0297
223.7978
226.9778
234.8579
238.8902
244.8871
248.9594
Relative Abundance
m/z

MS of Rbo5P.

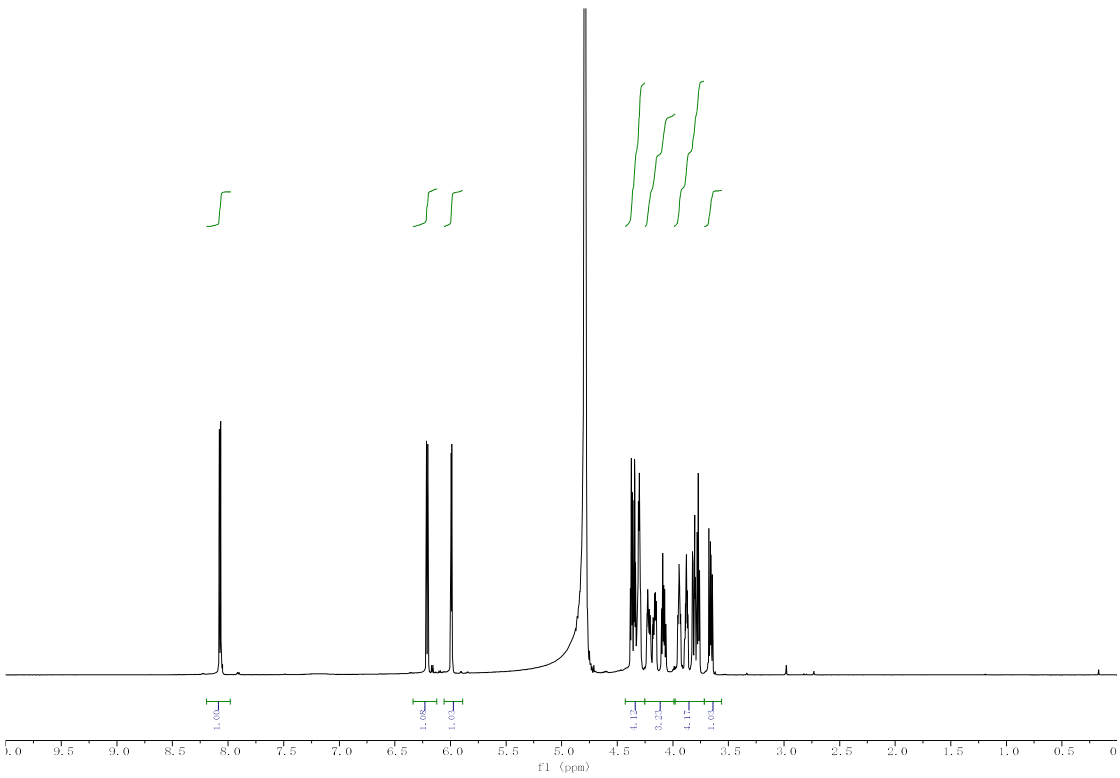


$^{1}H$ NMR of CDP-Rbo.

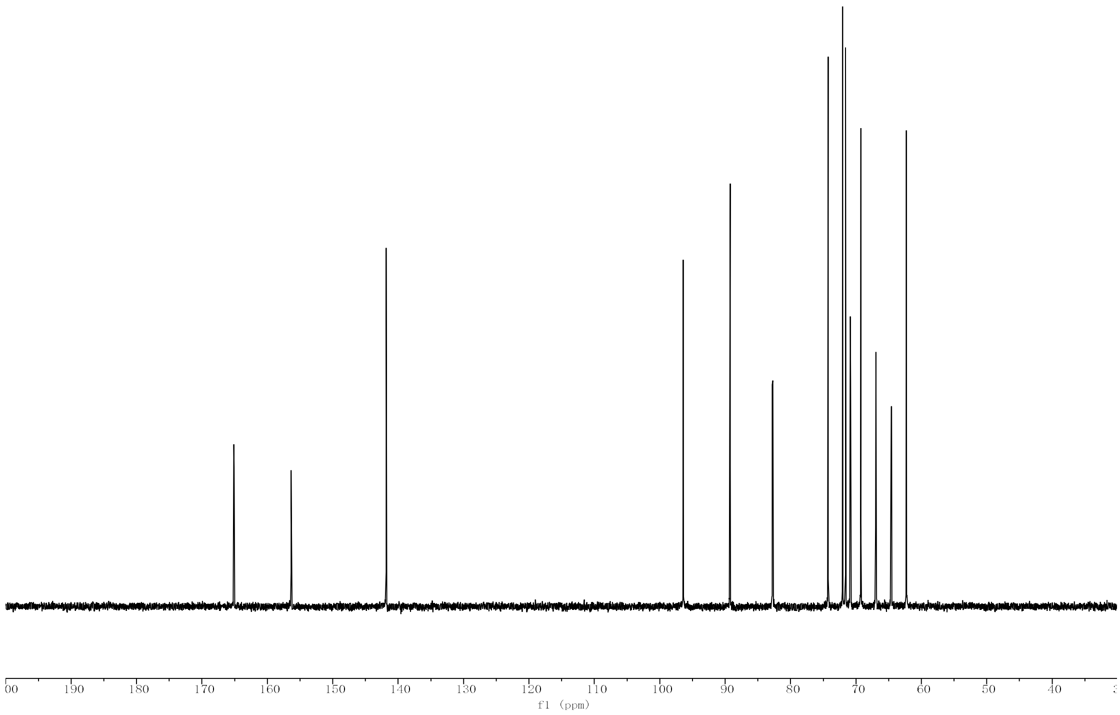


$^{13}C$ NMR of CDP-Rbo.

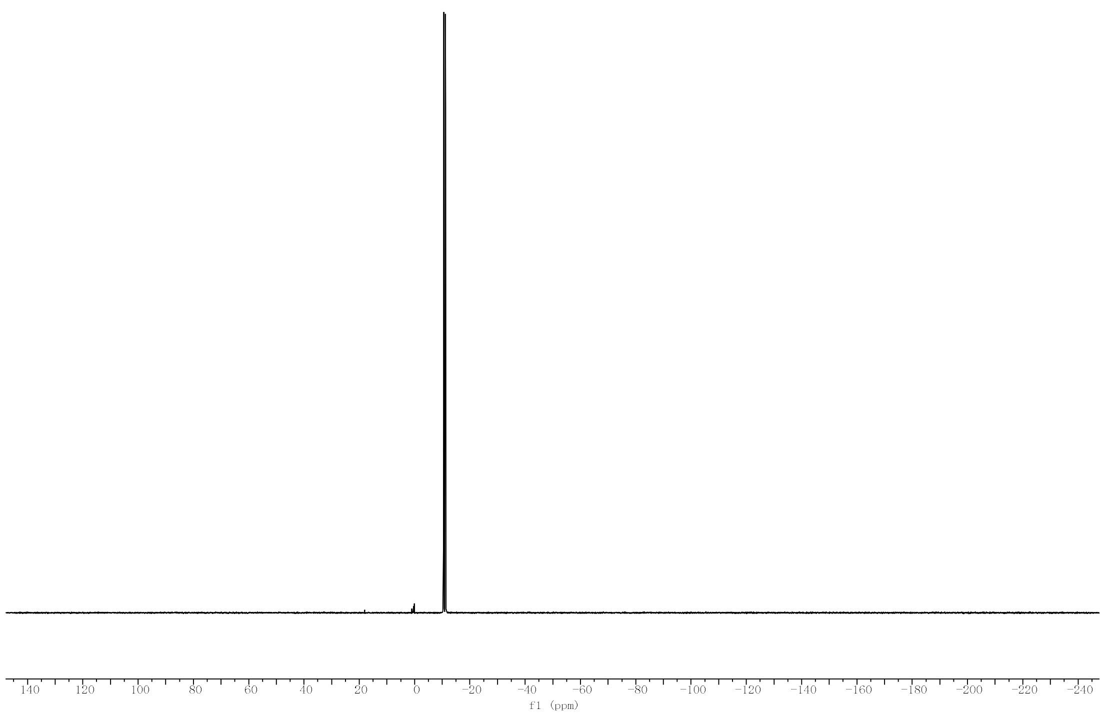


$^{31}$P NMR of CDP-Rbo.

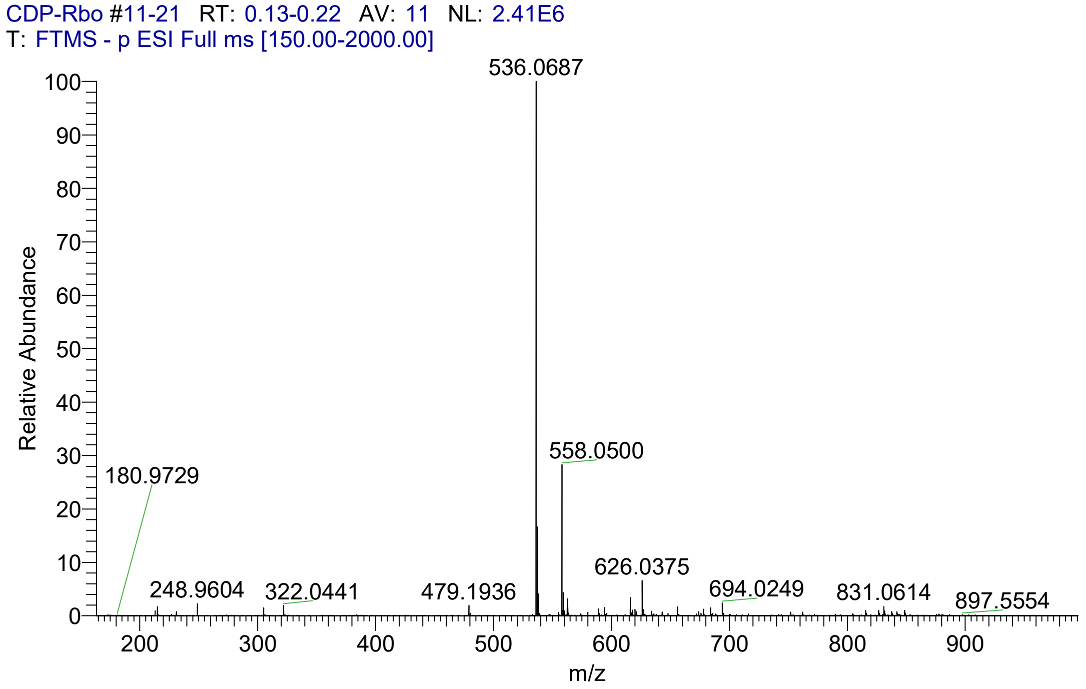


MS of CDP-Rbo.

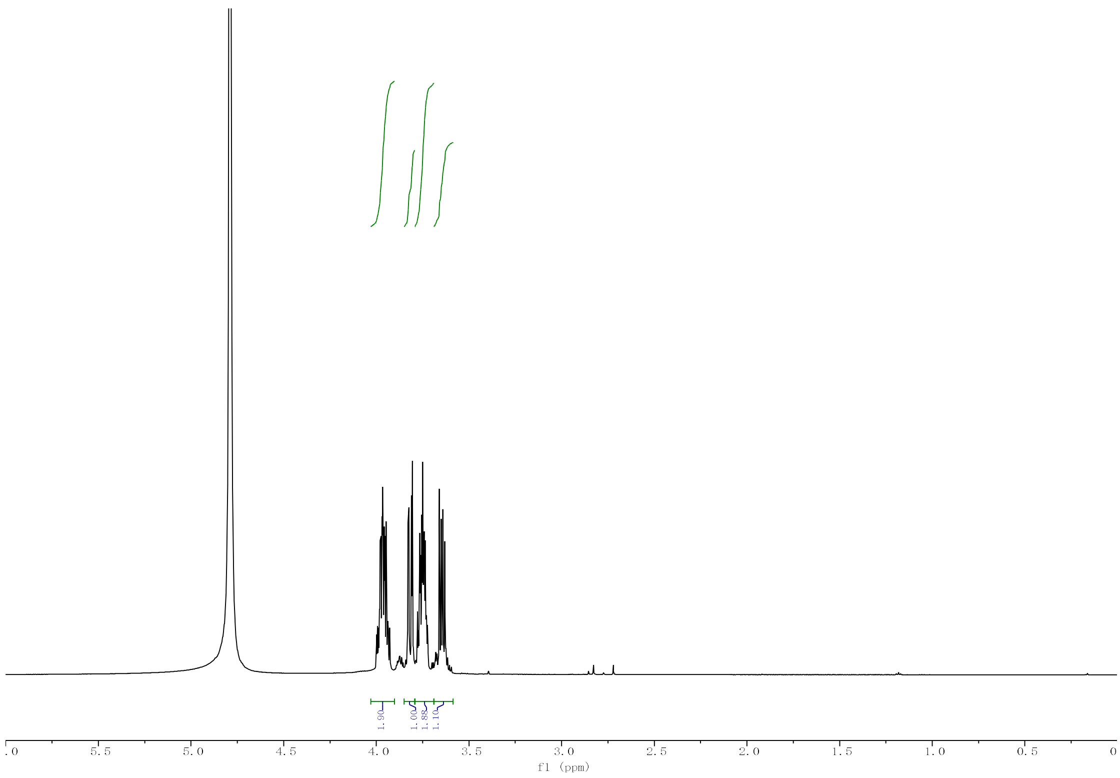


$^{1}H$ NMR of Erythritol-4P.

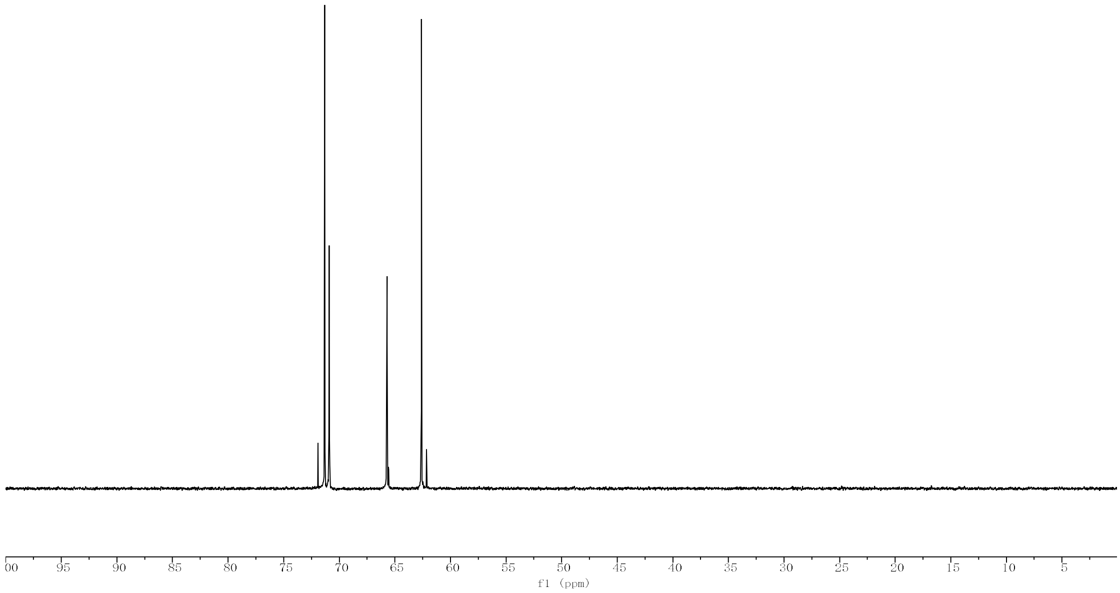


$^{13}C$ NMR of Erythritol-4P.

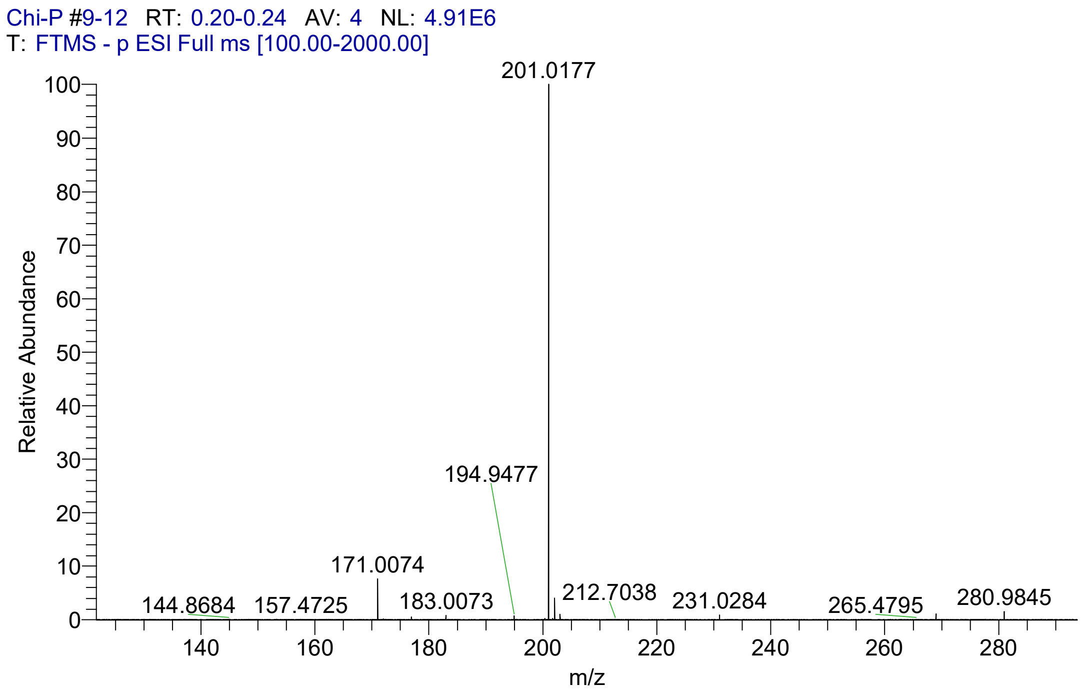


MS of Erythritol-4P.

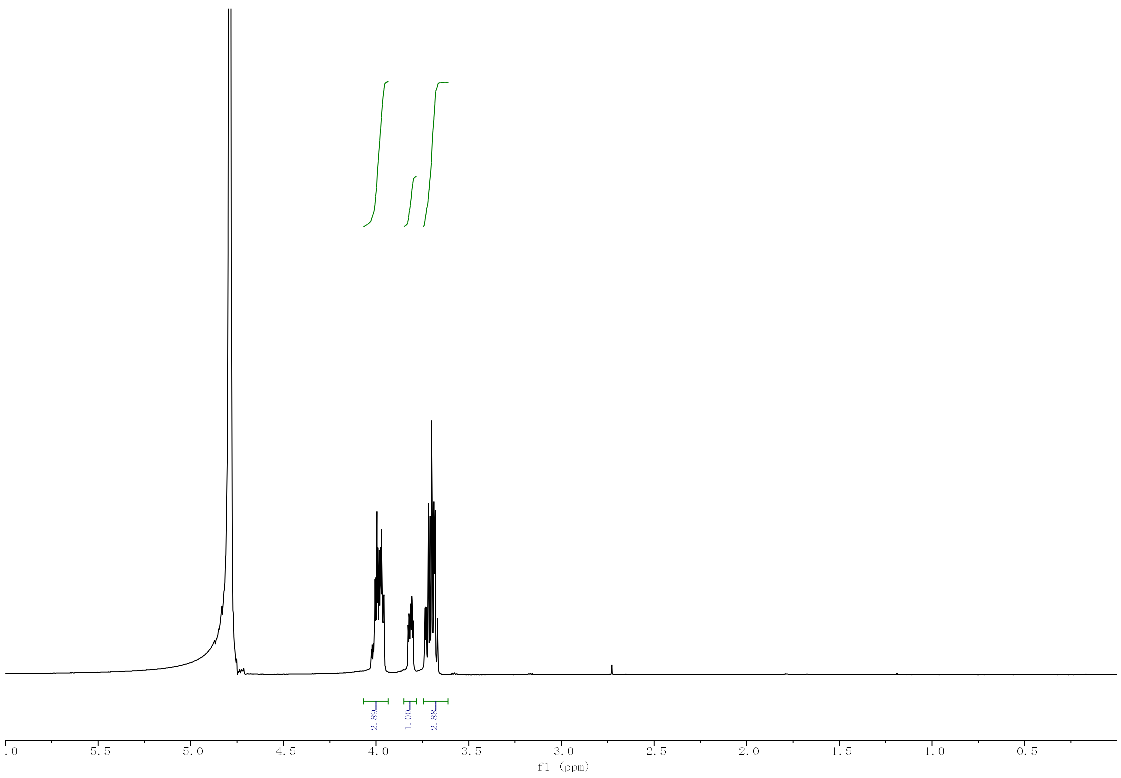


[1]H NMR of L-arabitol-5P.

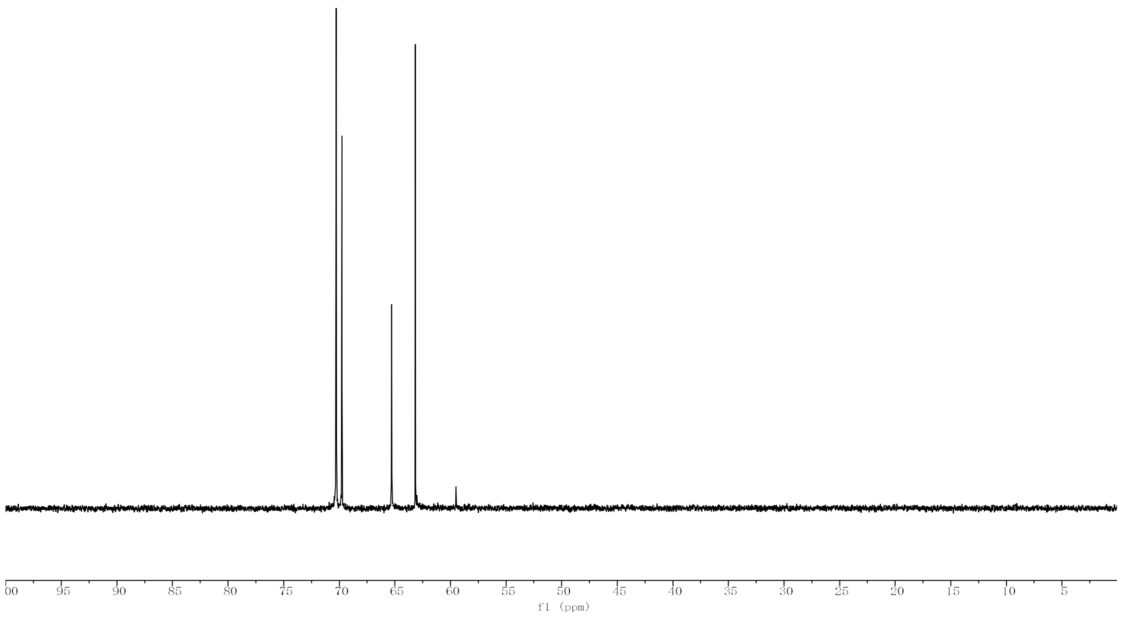


[13]C NMR of L-arabitol-5P.

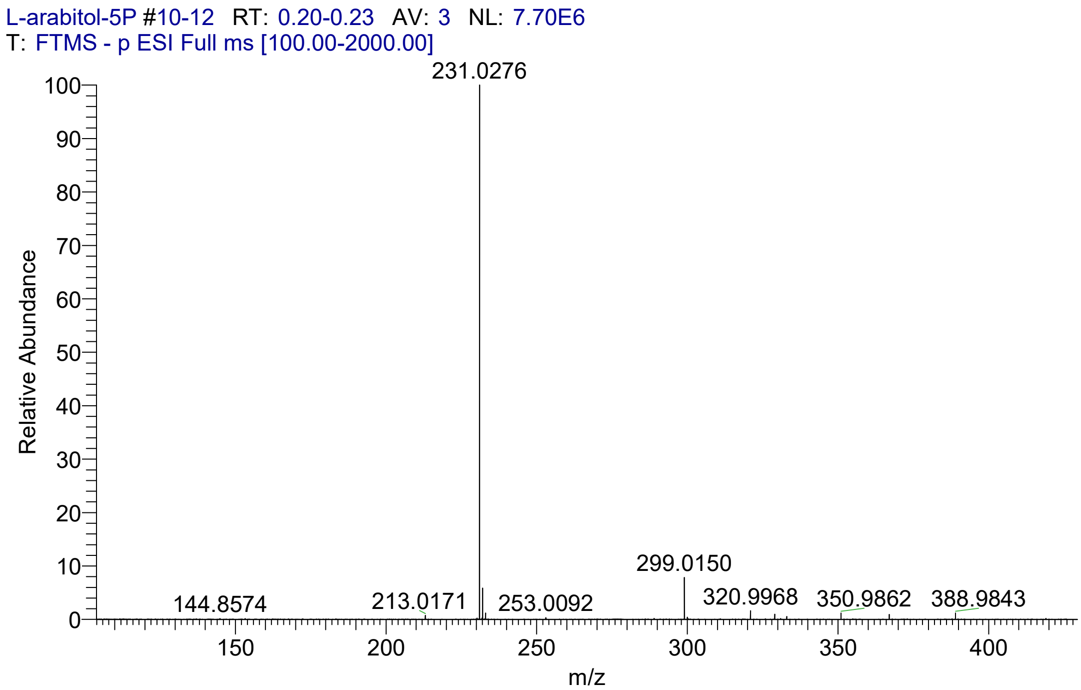


MS of L-arabitol-5P.

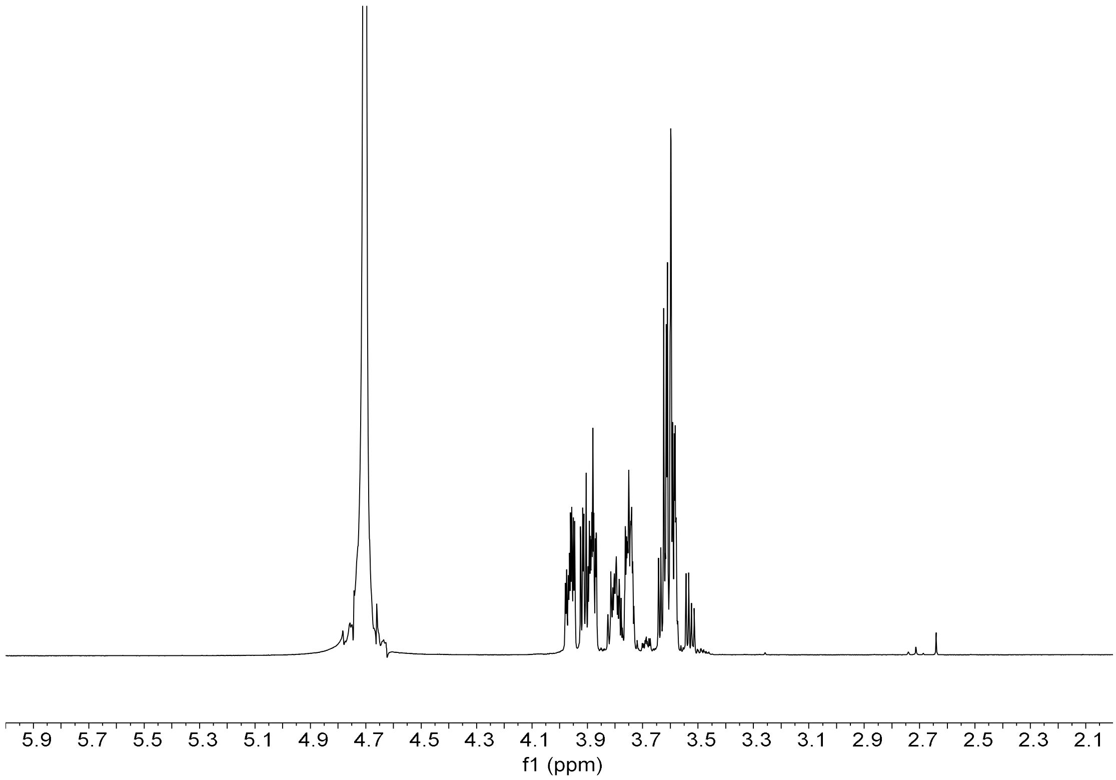


1H NMR of D-arabitol-5P.

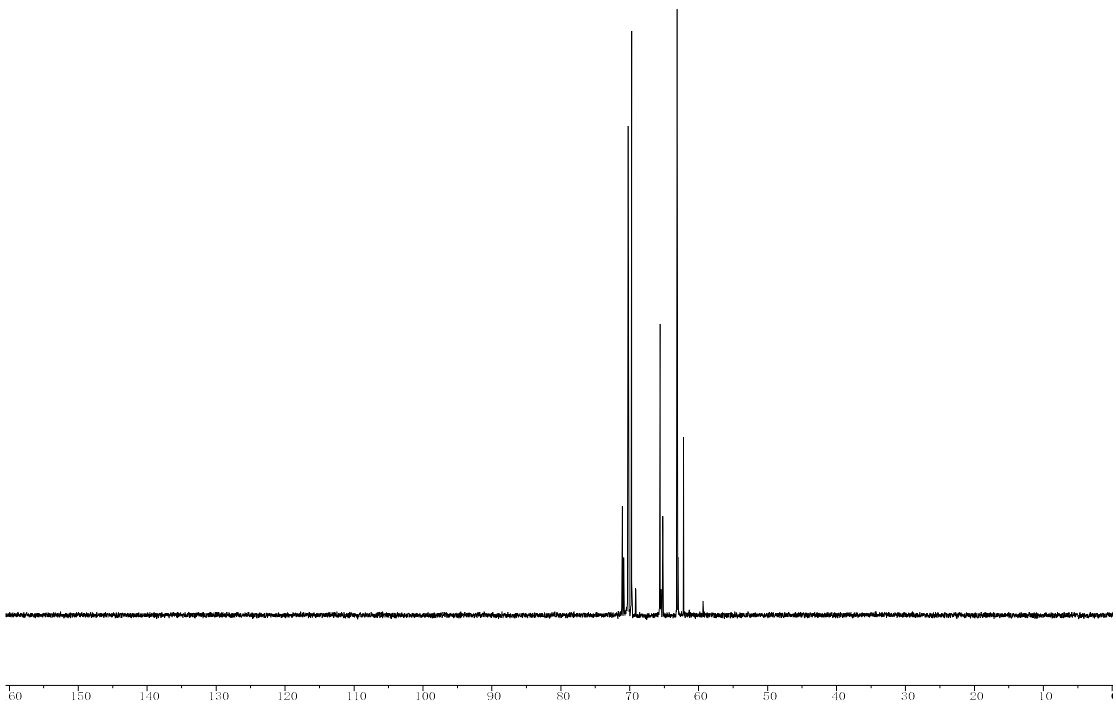


$^{13}$C NMR of D-arabitol-5P.

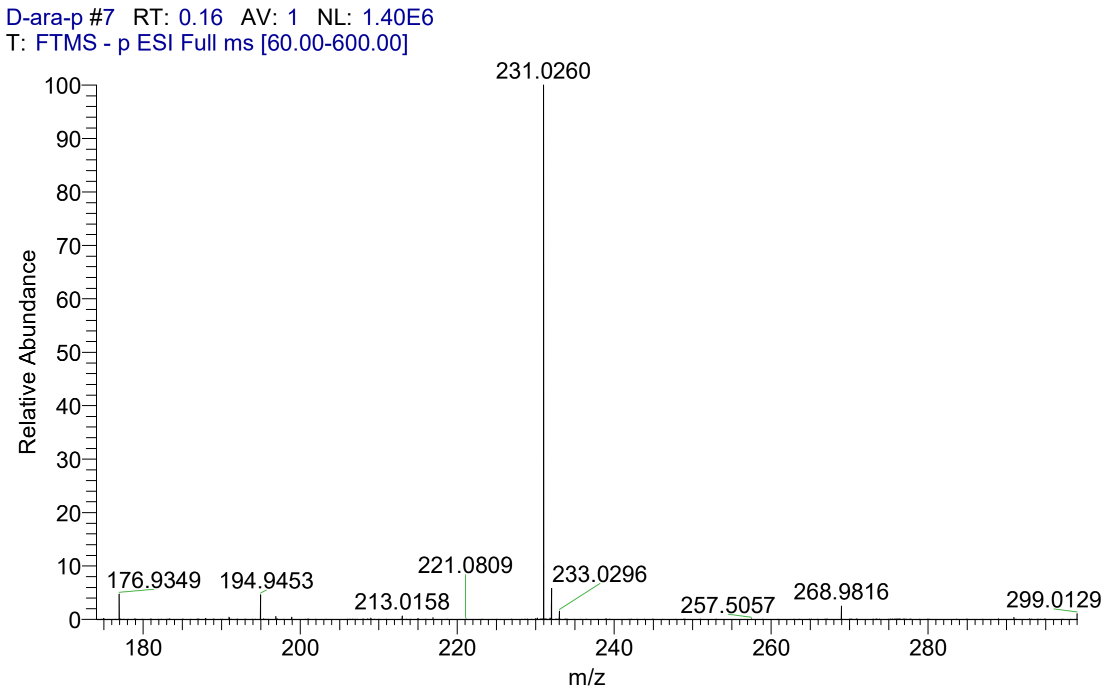


MS of D-arabitol-5P.

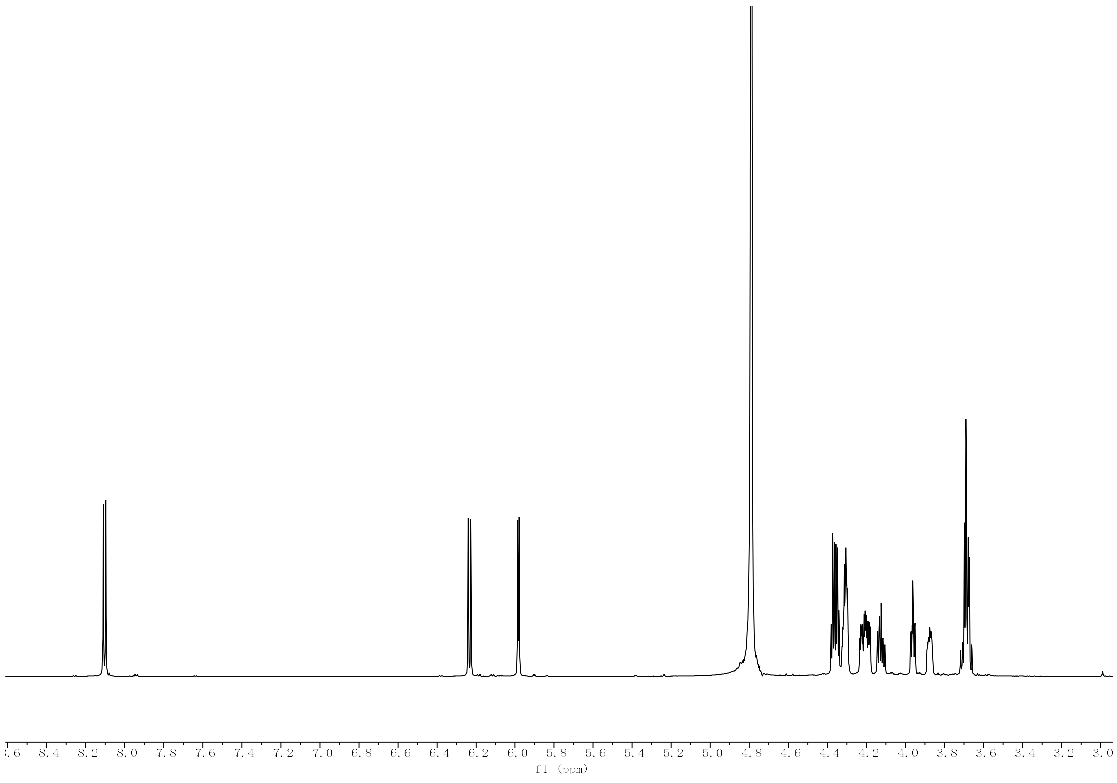


$^{1}H$ NMR of CDP-D-arabitol.

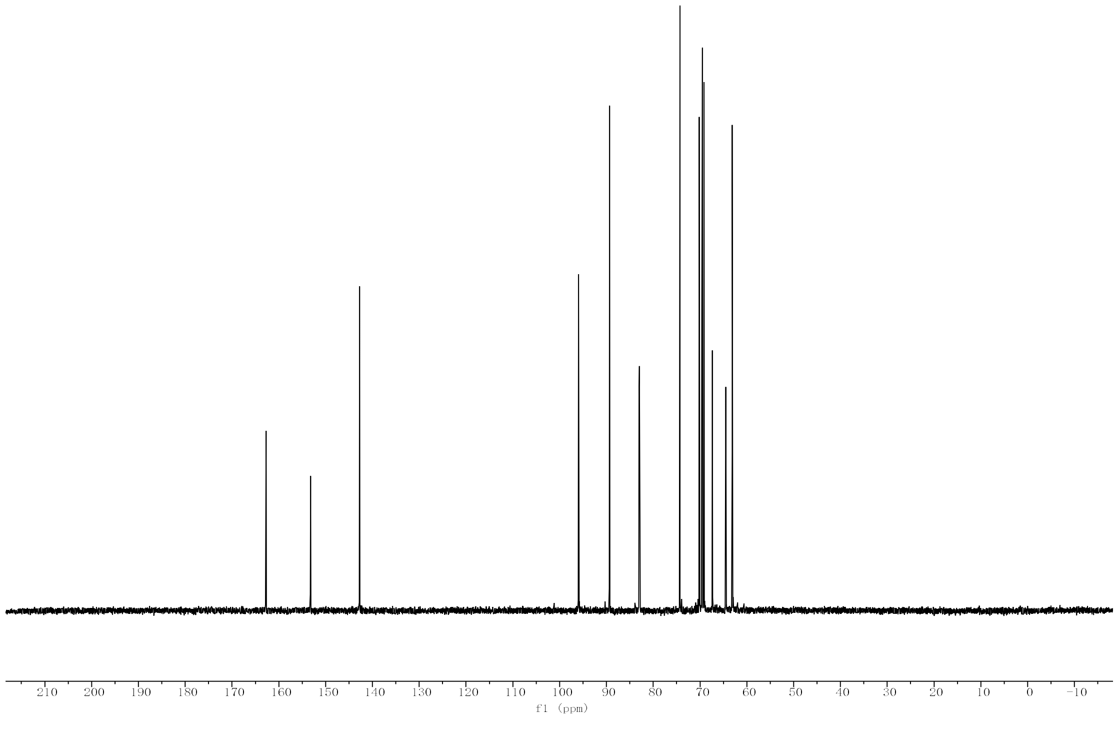


$^{13}C$ NMR of CDP-D-arabitol.

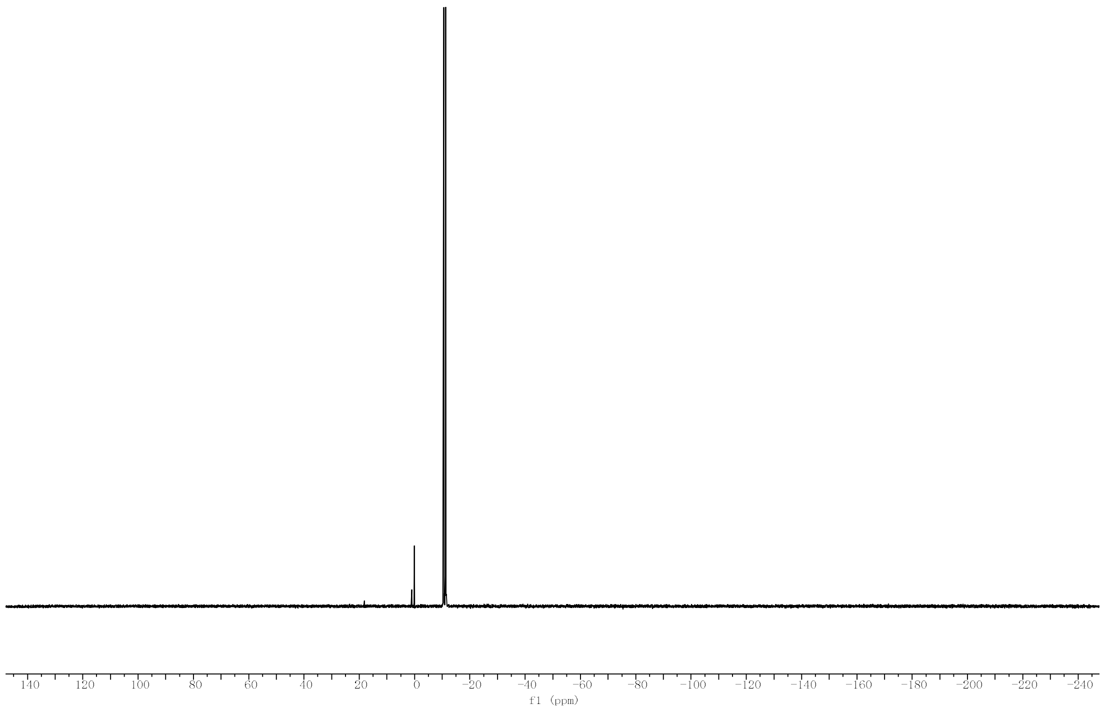


$^{31}P$ NMR of CDP-D-arabitol.

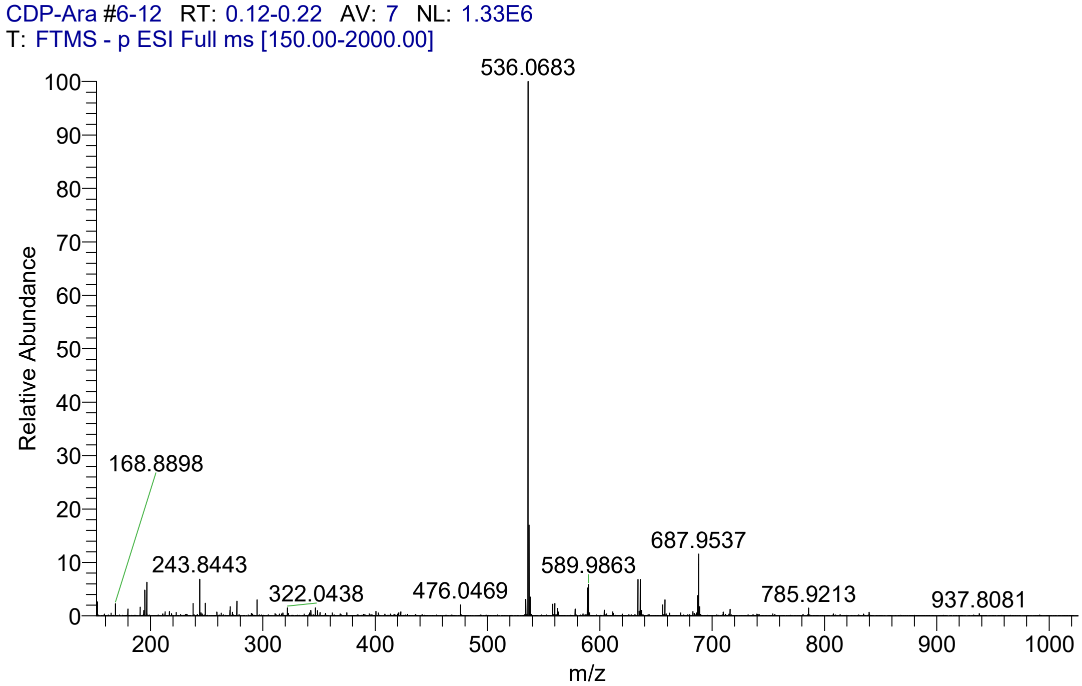


MS of CDP-D-arabitol.
